# *TimberTracer*: A Comprehensive Framework for the Evaluation of Carbon Sequestration by Forest Management and Substitution of Harvested Wood Products

**DOI:** 10.1101/2024.01.24.576985

**Authors:** I. Boukhris, A. Collalti, S. Lahssini, D. Dalmonech, F. Nakhle, R. Testolin, M. V. Chiriacò, M. Santini, R. Valentini

## Abstract

**Background:** Harvested wood products (HWPs) have a pivotal role in climate change mitigation, a recognition solidified in many Nationally Determined Contributions (NDCs) under the Paris Agreement. Integrating HWPs’ greenhouse gas (GHG) emissions and removals into accounting requirements relies on typical decision-oriented tools known as wood product models (WPMs). The study introduces the *TimberTracer* (TT) framework, designed to simulate HWP carbon stock, substitution effects, and emissions from wood decay and bioenergy.

**Results:** Coupled with the 3D-CMCC-FEM forest growth model, *TimberTracer* was applied to Laricio Pine (*Pinus nigra* subsp. *laricio*) in Italy’s Bonis watershed, evaluating three forest management practices (clearcut, selective thinning, and shelterwood) and four wood-use scenarios (business as usual, increased recycling rate, extended average lifespan, and a simultaneous increase in both the recycling rate and the average lifespan) over a 140-year planning horizon, to assess the overall carbon balance of HWPs. Furthermore, this study evaluates the consequences of disregarding landfill methane emissions and relying on static substitution factors, assessing their impact on the mitigation potential of various options. This investigation, covering HWPs stock, carbon (C) emissions, and the substitution effect, revealed that selective thinning emerged as the optimal forest management scenario. In addition, a simultaneous 10% increase in both the recycling rate and half-life, under the so-called “sustainability” scenario, proved to be the optimal wood-use strategy. Finally, the analysis shows that failing to account for landfill methane emissions and the use of dynamic substitution can significantly overestimate the mitigation potential of various forest management and wood-use options, which underscores the critical importance of a comprehensive accounting in climate mitigation strategies involving HWPs.

**Conclusions:** Our study highlights the critical role of harvested wood products (HWPs) in climate change mitigation, as endorsed by multiple Nationally Determined Contributions (NDCs) under the Paris Agreement. Utilizing the *TimberTracer* framework coupled with the 3D-CMCC-FEM forest growth model, we identified selective thinning as the optimal forest management practice. Additionally, enhancing recycling rates and extending product lifespan effectively bolstered the carbon balance. Moreover, this study emphasizes the necessity of accounting for landfill methane emissions and dynamic product substitution, as failing to do so may significantly overestimate the mitigation potential of implemented projects. These findings offer actionable insights to optimize forest management strategies and advance climate change mitigation efforts.

## 1. Background

Terrestrial ecosystems play a major role in the global carbon cycle owing to their inherent ability to absorb carbon (C) through photosynthesis and release it through respiration. The terrestrial biosphere provided a net sink for ~21% of carbon dioxide emitted by fossil fuel burning during the 1990-2021 period (1), with the major part occurring in forests (2). This climate change mitigation role of forests has been widely recognized by the United Nations Framework Convention on Climate Change (UNFCCC) being part of the periodical national greenhouse gas (GHG) inventories well as of many NDCs to the Paris Agreement (79% of the submitted NDCs covers the forest sector under mitigation targets, according to (3)).

Sustainably managed forests offer a dual avenue for GHG mitigation through processes that are mutually exclusive, namely sequestration and substitution (4). Reducing harvest in managed forests may yield to a positive impact on the forest C stock in the short to medium term but would adversely induce a long-term negative impact on the wood-chain value and a counterproductive effect specifically on carbon sequestration as aging trees exhibit decreased growth and carbon use efficiency (5). Conversely, the promotion of wood use would lead to the substitution of energy-intensive materials (e.g., steel or concrete) or fossil fuels, further compounded with the storage of carbon within harvested wood products (HWPs) (7). Given the trade-offs among different options (i.e., carbon sequestration, energy substitution, and material substitution), the most effective mitigation strategy would be the one that optimally balances and integrates all the mitigation components (8–10).

The acknowledgement of HWPs as integral to climate change mitigation occurs in many NDCs submitted by Parties under the Paris Agreement (3,11), within which countries voluntarily set binding GHG accounting obligations and targets to reduce GHG emissions and increase C-removals (12). The Intergovernmental Panel on Climate Change (IPCC) provides several approaches for estimating the GHG emissions and removals associated with HWPs and encourages deviating from the conventional ‘instantaneous oxidation’ approach (i.e., carbon in harvested biomass is considered as released into the atmosphere immediately after the harvesting) towards more accurate methods (13,14). Understanding the carbon dynamics of HWPs is critical for improving future forest-based climate mitigation strategies. While large-scale models such as CBM-CFS3 and EFISCEN are commonly used for national reporting under NDCs and GHG inventories, they primarily focus on forest carbon dynamics and provide only simplified representations of HWP carbon flows. In contrast, stand-level wood product models (WPM) offer detailed insights into carbon stocks and fluxes at the local scale, which are essential for refining forest management practices. The WPM developed in this study is specifically designed for stand-level applications rather than broader-scale reporting but can contribute to improving methodological approaches for tracking carbon storage in HWPs at fine spatial scales.

To accurately project storage and emissions in/from HWPs, models should incorporate all key components that influence C pools and emissions. However, most WPMs rarely account for all structural components, with aspects such as recycling, substitution, bucking allocation, and landfill gas emissions, specifically, often being overlooked (15,16). The practice of cascading wood products extends their lifespan, consequently delaying the release of GHG emissions into the atmosphere. A study conducted by Brunet-Navarro et al. using theoretical simulations to assess the impact of elevating recycling rates on carbon storage within wood products in the European (EU-28) wood sector and revealed compelling results. Specifically, an increase in the recycling rate from 10% to 20.9% between 2017 and 2030 was projected to yield a notable emission saving of nearly 5 MtCO_2_ (17). The substitution of materials with high energy requirements for production or the replacement of fossil fuels with less energy-demanding wood can permanently and cumulatively avoid emissions, provided that there is no leakage to other cross-sectors or replacement losses (18). For instance, in a case study comparing two functionally equivalent buildings – one constructed with a wooden frame and the other with a reinforced concrete frame – the manufacturing process emitted 45% less carbon in the wooden structure while also requiring less energy (19), which underscores the importance of considering the substitution effect in the WPMs design. A critical point regarding product substitution is that most analyses assume a constant displacement factor value over time, even though this value may decrease due to factors such as evolving manufacturing methods, improvements in energy efficiency, and the redesign of the energy mix (18,20). Recent studies examining the fundamental assumptions underlying the projection of mitigation benefits related to substitution, including the static nature of displacement factors, have found that this substitution effect has been overestimated by 20% to 96% (21). It is thus essential that future model implementations account for these potential variations. From another perspective, the bucking allocation involves the disaggregation of logs into different HWPs based on quality and dimensional criteria. It is crucial to consider this process when assessing the effects of management (e.g., rotation, thinning intensity or interval). For instance, studies that include the bucking allocation process suggest considering longer rotations for optimizing carbon stock in the forest sector (e.g., see (8)), while those excluding it recommend shorter rotations (e.g., see (22)). This is because models not incorporating the bucking allocation module use predefined default values to allocate harvested volume to HWPs. In principle, the higher the productivity, the greater the quantity of products – a scenario typically observed in shorter forest rotations. In contrast, when the bucking allocation is accounted for, the optimization of C stock would favor the production of HWPs with a longer lifespan, generally derived from larger tree stems – a scenario typically observed in longer rotations modeling analyses (23). In landfills, the limited availability of oxygen creates unique conditions for the decomposition of major wood polymers. Under anaerobic conditions, microbial activity breaks down a portion of the dissolved organic carbon (DOC) into methane (CH_4_) and carbon dioxide (CO_2_). While many models that include landfills primarily estimate C stock, they often overlook the varying global warming potential of different landfill gases. As a result, solely assessing changes in carbon stock within landfills is insufficient for accurately estimating total greenhouse gas (GHG) emissions.

In this paper, we have pioneered the development of an open-source Python-based model, named *TimberTracer (TT)*, to comprehensively account for the various components that directly or indirectly influence C pools and emissions – some of which are typically overlooked by existing forest/vegetation models – and to provide insights into the temporal dynamics of both carbon storage and emissions associated with HWPs. To assess the climate change mitigation potential of the entire forest sector, the model can be seamlessly interface with any forest growth model, whether individual-tree or stand-based level, such, as e.g., GO+ (24) or 3D-CMCC-FEM (25). The main objective of this study is to examine the impact of forest management and wood utilization on the mitigation potential of HWPs. Focusing on the *Pinus nigra* subsp. *laricio* (Poiret) forest located in the experimental Bonis watershed in southern Italy (26), we employ a carbon modeling framework by combining the 3D-CMCC-FEM with *TT*. This integrated framework is used to simulate the evolution over 140-years of the HWP stock, carbon emissions, and the substitution effect under three forest management schemes and four wood-use scenarios, as elaborated in subsequent sections. Furthermore, to determine the influence of dynamic substitution and methane release from landfills on the mitigation potential of various forest management and wood-use scenarios, we use two different model implementations: one that includes methane emissions from landfills and dynamic product substitution, and another that models solely the carbon stock changes in landfills while considering static product substitution.

## 2. Methods

### 2.1. Wood products modeling, an overview

WPMs are implemented diversely based on their scopes and objectives. An integral model should include –in principle– these six following components: (i) *bucking allocation,* which involves the disaggregation of stems into different logs destignated to the production of differentiated products according to a set of dimensional and quality criteria typically established by wood industry professionals. Considering this component as an integral part of the TT model, rather than relying on a priori allocation factors, is crucial for minimizing errors resulting from simplification; (ii) *industrial processes,* that involve the transformation of raw wood into finished or semi-finished products, as well as the recycling or disposal of products when reaching the end of their use. Those processes are characterized by a set of allocation parameters either derived from expert knowledge, local surveys, or previous studies; (iii) *carbon pools,* which refer to the reservoirs of carbon stored in the diverse wood products currently in use or at disposal sites. HWPs are characterized by an average time in use, primarily linked to their intended purpose. For instance, construction wood is typically long-lived compared to pulpwood, given its use in applications that demand high durability and longevity. In contrast, pulpwood is primarily utilized for paper and pulp production, and its intended use does not require the same level of longevity; (iv) *product removal,* that refers to the point in time when products are retired from use. To estimate the product removal rate (also known as ‘retirement rate’), C-retention curves are used. They are based on the cumulative function of a chosen statistical distribution (e.g., Weibull, uniform, linear, and normal distributions) defined by one or more of the following parameters, e.g., the time when 50% of the initial C-stock is left (also known as half-life time), the time when 5% of the initial C-stock is left, the average life, and the maximum retirement rate; (v) *recycling,* that involves the transformation of HWPs after reaching the end of their use into new products. This theoretically induces a reduction in the retirement rate, as an additional amount of products is reinjected after each projection compared to the scenario when recycling is not considered; and (vi) *substitution,* which refers to the displacement effect resulting from the use of wood to substitute functionally equivalent energy-intensive materials or fossil fuels. The major components and processes described above are graphically illustrated in **Figure 1**.

**Figure 1:**
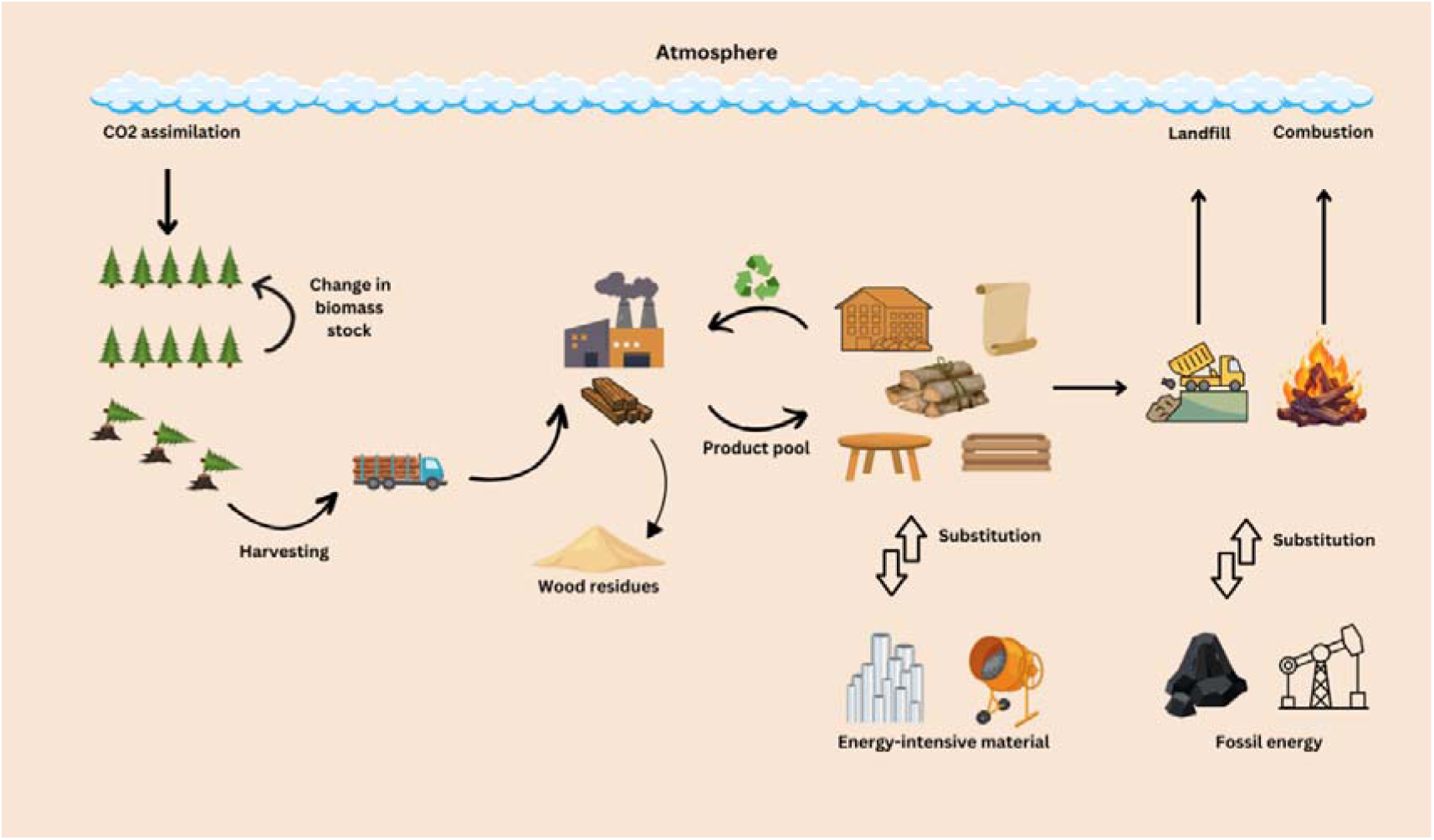
Lifecycle of HWPs - Illustration depicting key stages from production through utilization to end-of-life and natural decay.

### 2.2. Model description

*TimberTracer* (TT), a WPM based on the material flow method, was implemented using Python as its programming language. The model is specifically tailored for a comprehensive analysis of wood products, and it encapsulates a robust framework for carbon storage analysis, enabling users to meticulously evaluate the quantity of carbon stored within diverse wood products. Furthermore, TT incorporates temporal insights, enabling stakeholders to scrutinize the evolving patterns of carbon storage and emissions over time. Remarkably, to assess the climate change mitigation potential for the entire forest sector, TT can seamlessly interface with theoretically any forest growth model, whether individual-tree or stand-based level, such, as GO+ (24) or 3D-CMCC-FEM (25).

By incorporating all the previously described components (see previous **section 2.1**), TT comprehensively accounts for and simulates the temporal dynamics of GHG emissions and removal across all carbon pools outside the forest, including changes in HWPs and disposal sites. Furthermore, the model considers the substitution effect and captures the temporal dynamics of material and energy substitutions. Additionally, TT offers the flexibility of utilizing both tree- and stand-level inputs, facilitated by its integration of a stand structure generator, enabling the transition from stand state descriptors to tree state descriptors. The TT model offers the ability to simulate the entire carbon accounting process using the function ‘`run_model()’`. Additionally, it is designed to be modular, allowing running various simulations independently, at specific stages of the C accounting process, as described in **Figure 2**. Hereafter, we explicitly introduce each module, outlining its role and the functions it encompasses.

**Figure 2:**
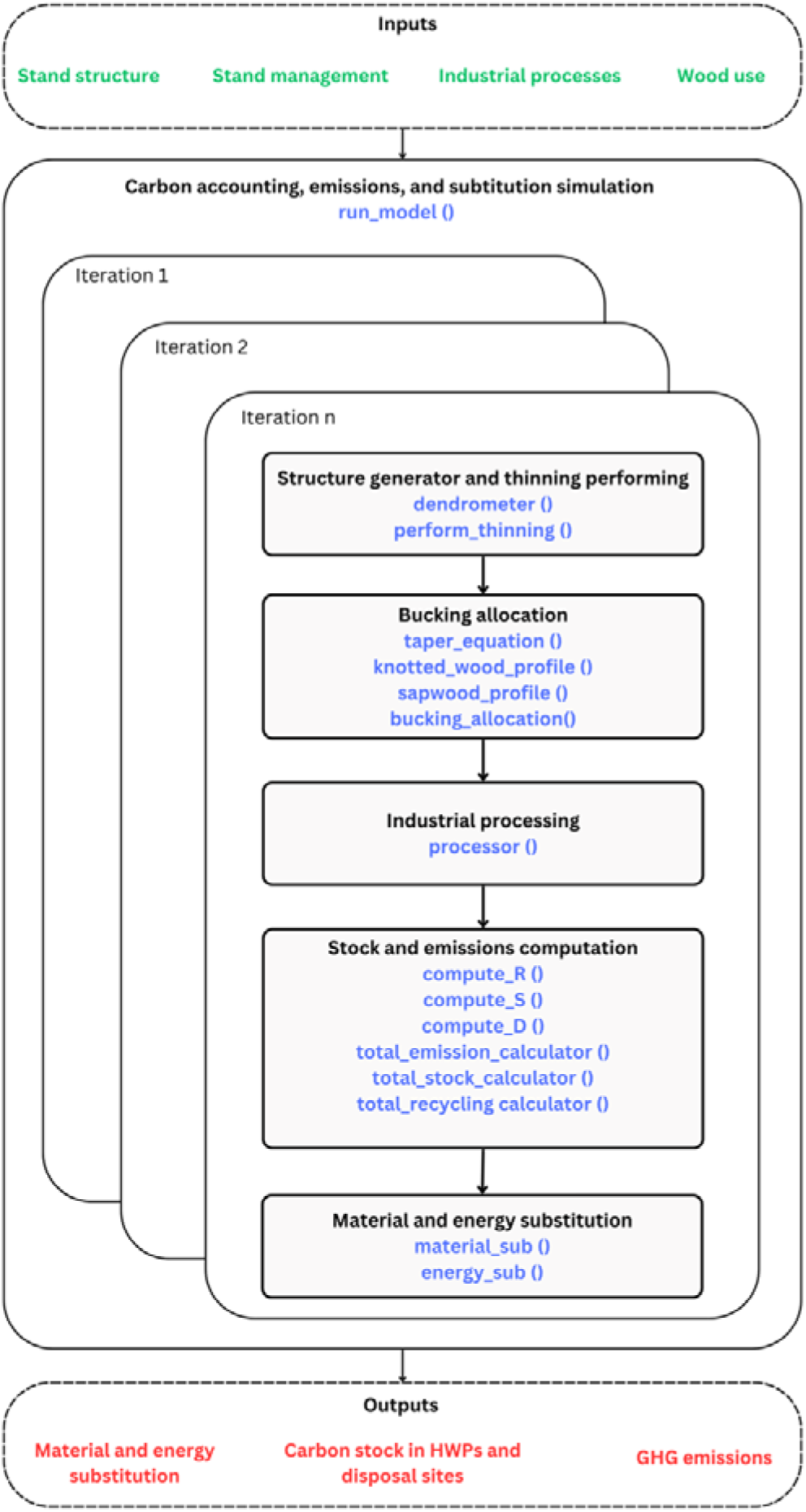
Flowchart of the TT model. In green the inputs, in red the outputs, in black the modules, and in blue the functions within the modules. The number of iterations is equal to the number of management interventions.

### 2.3. Model inputs and requirements

The TT model requires a set of input data for its initialization, including: i- stand structural data (DBH, height, stand density, basal area, bark thickness, sapwood, and heartwood areas); ii- forest management data (planning horizon, rotation period, rotation number, thinning age, thinning intensity, thinning nature); iii- industrial process data (product, priority, number, log length, log diameter, quality criteria, process efficiency); iv- wood use data (lifespan, recycling rate, reallocation scheme, displacement factors). Furthermore, TT necessitates the specification of a well-defined set of parameters mainly required by the dendrometer and the bucking allocation module (for further information on the model parameters and data inputs, please refer to the table in **Supplementary 3**).

### 2.4. Model outputs

The ‘`run_model()’` function serves as an integrated coordinator, orchestrating the seamless execution of independently developed modules introduced earlier to generate a comprehensive output. The model can be run for a planning horizon (PH) that may encompass multiple rotations (R). The outputs provided include the simulations of carbon stock in HWPs, the annual and cumulative emissions, the annual and cumulative material and energy substitution, and the yearly recycling (all expressed in tC ha^−1^).

### 2.5. Model structure

The TT model is designed to be modular. In this section, we will introduce each module individually, explaining the theory and objectives of each (for further information refer to **Supplementary 1**).

#### 2.5.1. The structure generator and thinning performing module

TT is designed for compatibility with both tree- and stand-level data, providing, to some degree, flexibility in data input. This capability is realized through the incorporation of a structure generator module that implements the statistical distribution (e.g., Weibull distribution) of the diameter at breast height (DBH) within the stand level. It also assigns trees to different bio-sociological classes of status (i.e., dominant, co-dominant, intermediate, and overshadow trees) based on their total tree height, as implemented in the ‘`dendrometer()’` function. TT further considers thinning operations, characterized in the model by three descriptors: type, intensity, and timing, implemented in the ‘`perform_thinning()’` function, which renders dendrometry information about each individual thinned tree.

##### 2.5.1.1. The structure generator

The transition from stand state descriptors to tree state descriptors would be feasible if the stand structure is well understood. The latter is typically likened to a statistical distribution. There are numerous mathematical formulations for statistical distributions in forestry, with one of the most employed being the Weibull distribution (28). The probability density function of a Weibull random variable denoted as *X* ~ *Weibull*(*sh*, *sc*, *loc*) is:

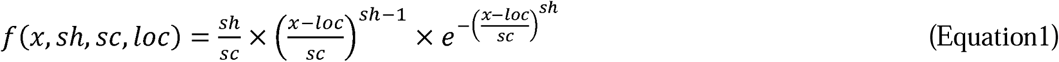

Where *f* is the probability density function, *x* is the random variable, *sh* represents the shape, *sc* denotes the scale, and *loc* indicates the location.

If the location is not provided, but the shape and scale are (2-parameter Weibull distribution), then the location will be predicted using the following equation:

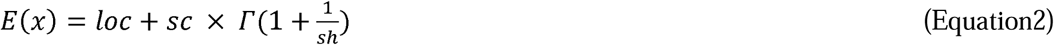

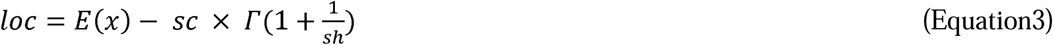

Where *E(x)* is the mathematical expectancy of *x* and *Г* is the gamma function.

After fitting the parameters of the Weibull distribution supposed to represent the structure of the forest stand at different stages of development, the next step is to compute the number of trees by each diameter class. For this specific purpose, the cumulative distribution function of the Weibull function is applied to the concerned diameter class following the equation:

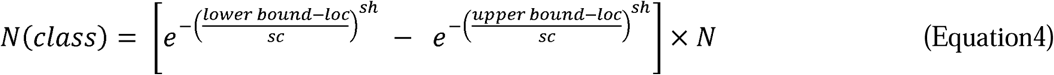

Where *N* represents stand density (ha^-1^), and the lower bound and upper bound define the DBH class interval.

##### 2.5.1.2. The thinning process

A thinning operation involves removing tree stems to benefit a tree or a group of trees deemed essential to ensure optimal growth conditions (29). It focuses on enhancing wood quality and stand stability. The thinning intervention can be characterized by three elements: (i) the type of cut (from the above, from the bottom, or neutral), which denotes the distribution of removed stems among various diameter categories of standing trees within the stand; (ii) the thinning intensity which expresses the magnitude of the extraction conducted within the stand. It can be quantified by both the number of stems harvested and the implementation rate relative to the before-thinning stand population; (iii) the period, commonly referred to as rotation, is the time interval between two successive thinning events within the stand. It varies depending on the tree species, age, and site conditions (30).

Bio-sociological tree status plays an important role in some thinning concepts, for example, for the selection of thinning trees in thinning from above and below (31). A simplified classification groups trees into four categories: i- dominant trees; ii- co-dominant trees; iii- intermediate trees; and iv- overshadow trees. This classification consists of the definition of the position of each tree in terms of its height by reference to the other trees of the stand. As part of the process, the dominant height of the plot (h_95%_) as well as the height to the base of the crown (hc) are calculated. The classification of the trees into a scale is performed following the rules presented in the **Table 1**.

**Table 1:** Bio-sociological status criteria based on the tree height.

| Criteria | Bio-sociological status |
| --- | --- |
| $hi \geq h_{95\%}$ | Dominant tree (1) |
| $\frac{h_{95\%} + hc}{2} \leq hi < h_{95\%}$ | Co-dominant tree (2) |
| $hc \leq hi < \frac{h_{95\%} + hc}{2}$ | Intermediate tree (3) |
| $hi < hc$ | Overshadow tree (4) |

The thinning operation follows the approach implemented by (32). For thinning from below and thinning from above, trees belonging to a specific bio-sociological class are prioritized for each thinning type. In the case of neutral thinning, trees are harvested without consideration for their bio-sociological status. Further details are provided in the following.

Thinning from below (4+3) => 2

The process consists of removing trees belonging to the dominated bio-sociological sub-groups. The process of removal is parallel in 4+3 (Overshadow and Intermediate). If the removal amount is not reached by 4+3, the process continues sequentially in sub-group 2 until reaching the required number of trees satisfying the initial condition.

Thinning from above (1+2) => 3

The process consists of removing trees belonging to the dominating bio-sociological sub-groups. The process of removal is parallel in 1+2 (Dominant and Co-dominant). If the removal amount is not reached by 1+2, the process continues sequentially in sub-group 3 until reaching the required number of trees satisfying the initial condition.

Neutral thinning (1+2+3+4)

The process consists of removing trees regardless of the bio-sociological sub-group to which they belong. The process is parallel in 1+2+3+4 and continues until satisfying the required removal amount.

#### 2.5.2. The bucking allocation module

TT allocates logs from thinned trees to HWPs, considering determinant factors such as stem log diameter and quality. The qualitative grading of logs involves several criteria, with the proportion of knotted wood and sapwood-to-heartwood ratio being among the most commonly used in the literature (33–35). The bucking allocation is practically implemented within the model through two sequential steps. The first step involves dressing the stem profile of each individual tree using the ‘`taper_equation()’` function which renders the diameter over bark (dob) at any point along the stem. Simultaneously, the knotted wood profile is dressed using the ‘`knotted_wood_profile()’` function while the sapwood and heartwood profiles are established with the ‘`sapwood_profile()’` function. The second step consists of combining the tree stem profile (including taper, knotted wood, sapwood, and heartwood) with the bucking allocation criteria defined by wood industry professionals to disaggregate trees into different logs. This entire process is implemented in the ‘`bucking_allocation()’` function (please refer to **Supplementary 2** for more information on the bucking allocation criteria).

##### 2.5.2.1. Stem profile generator

To achieve a precise stem profile dressing, the TT uses a comprehensive set of equations, which will be thoroughly described in this section (see **Figure 3** for a graphical representation of the stem profile).

**Figure 3:**
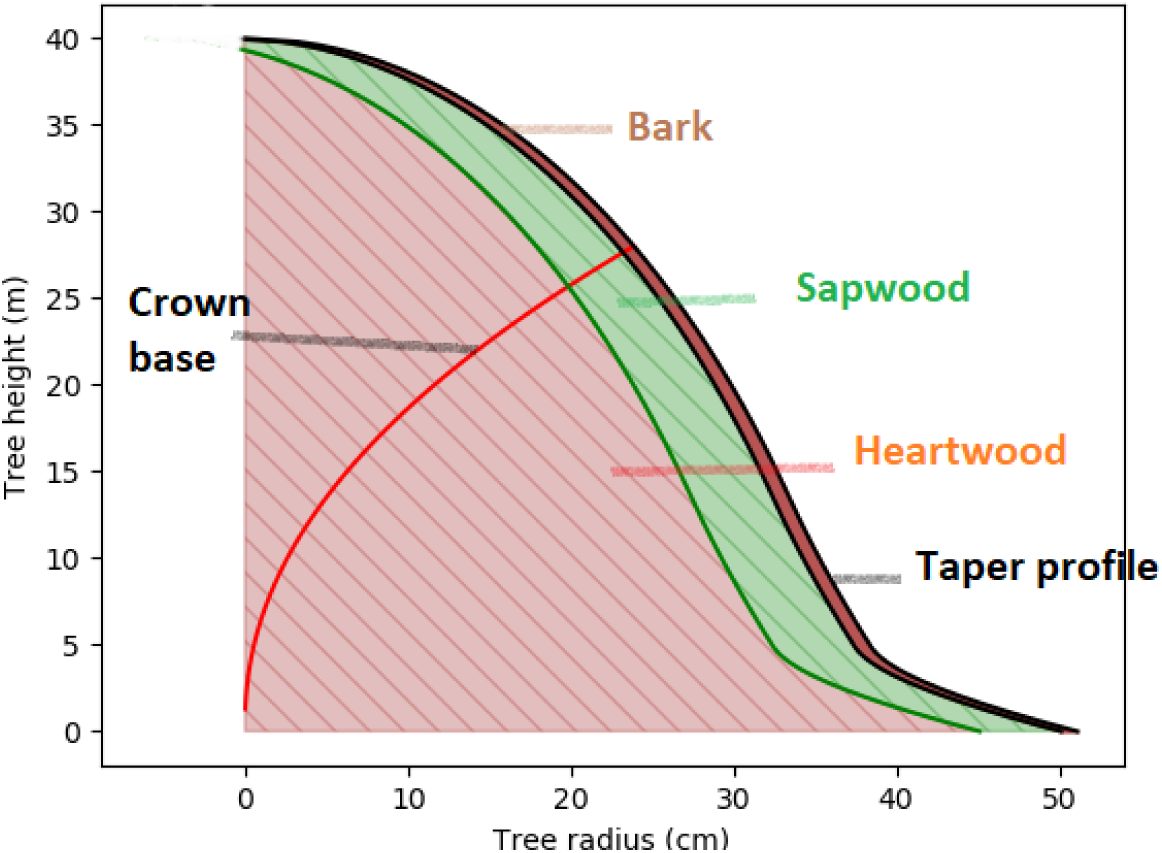
Graphical representation of stem longitudinal profile

The taper profile refers to the degree to which a tree’s stem diameter decreases as a function of height above ground. Taper is often represented by mathematical functions fitted to empirical data, called taper equations. One such function, attributed to (36) and used by default in the model is:

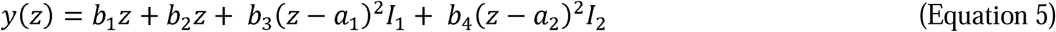

where 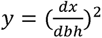; dx = is the upper stem diameter over bark (dob) at a given height h of the tree, 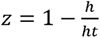 = the complement of the relative height with ht being the tree total height; a_1_and a_2_ = join points to be estimated from the data, *I_k_* = 1 if *z* > *a_i_* and 0 otherwise, *k* = 1,2, *b_p_’*s = regression coefficients with, *p* = 1, 2, 3, 4.

The crown base height (*h_c_*) refers to the level of insertion of the last branch of the crown within the stem, it is generally predicted from the total height using an allometric equation. One such equation is the power function expressed as follows:

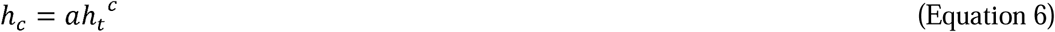

where *a* and *b* correspond respectively to the amplitude and the exponent of the relation.

The DBH of a tree could be also predicted from its total height. One such function is the power function expressed as follows:

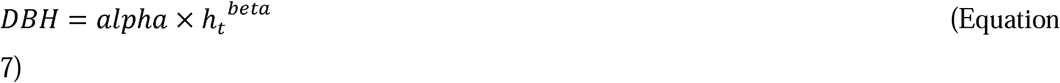

where *alpha* and *beta* correspond respectively to the amplitude and the exponent of the relation.

In tree analysis, a crucial step involves shaping the crown base profile to differentiate knotted from intact wood. To establish this profile, a combination of the three equations (5–7) is utilized. In practice, the approach entails reconstructing the historical crown base height limits and considering that the area from the curve towards the bark represents the knotted wood, while the area from the curve towards the pith represents the intact wood.

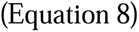

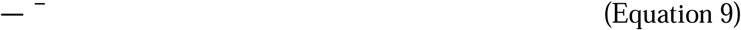

where represents the complement to the relative height at the level of the crown base and *d_c_* refers the diameter of the intact wood.

The sapwood profile is determined based on the simplified assumption that the sapwood width and bark thickness remain constant within the tree level (37). The width of knotted wood, which naturally varies within the tree level, is calculated as the difference between the diameter at a specific height obtained from the taper profile and the sum of the sapwood width and bark thickness.

##### 2.5.2.2. Stem disaggregation

Stems are disaggregated into different logs based on specific criteria, typically established by wood industry professionals. These criteria, tailored to each species, include both dimensions and quality considerations, and they are organized hierarchically. To elaborate, the model initially checks if the stem aligns with the criteria for producing a designated log intended for a specific product. If this is not the case or if the maximum desired number of this specific log has been reached, the model then advances to the next HWP based on a hierarchy defined by the user. In TT, log dimensions are characterized by the length and small end diameter of the log, while log quality is assessed through ratios of knotted wood (*Ø_KW_*) to heartwood (*Ø_HW_*) and knotted wood to small end diameter (*Ø_SE_*). The model checks these criteria (e.g., *Ø_SE_* ≥ 25 cm, (*Ø_KW_ / Ø_HW_*)^2^ ≤ 13%, and *Ø_KW_/ Ø_SE_*≤ 30%) by implementing a bisection algorithm that automatically performs the search for the corresponding height validating the given criteria.

#### 2.5.3. The processor module

Logs are processed to match their intended products, employing the standard industry method for each type of product. The efficiency rate of this industrial transformation, a theoretical measure, depends on how the processing is done, and this varies based on the targeted product. In *TT*, the production of HWPs is calculated using the efficiency rate which is product-specific. Subsequently, any process losses (i.e., 1 – efficiency) are redistributed among other products according to a scheme defined by the user. This process is implemented in the ‘`processor()’` function.

#### 2.5.4. The stock and flow module

TT simulates the evolution of carbon in HWPs and disposal sites including both mill and landfill sites throughout the planning horizon (PH) using the ‘`total_stock_calculator()’` function. After each annual projection, a portion of the carbon in HWPs undergoes retirement based on product-specific decay function (refer to ‘`compute_D()’` in the model). Subsequently, a fraction of the retirement is recycled and reinjected into the HWPs carbon pool (refer to ‘`compute_R()’` in the TT model), another portion is directed to firewood, while the remaining part is sent to landfills. Furthermore, TT simulates the evolution of the total emissions resulting from the firewood combustion and the decay of carbon pool in disposal sites, accounted for as an instantaneous oxidation, using the ‘`total_emission_calculator()’` function.

In TT, the product removal rate (i.e., retirement rate) is determined using the cumulative distribution function (CDF) of a normal distribution, which is defined by its mean and standard deviation. The mean represents the average lifespan of a product, while the standard deviation reflects the change in the dynamic retirement, typically expressed as a fraction of the average lifespan (e.g., a fraction of 1/3 as used in (17)). In the following, the CDF formulation is presented.

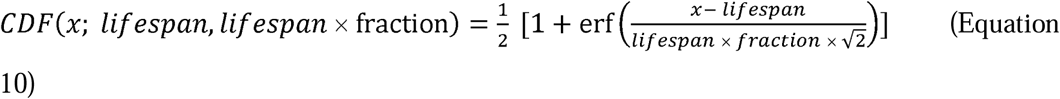

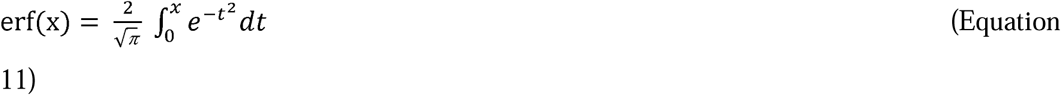

where CDF is the cumulative distribution function and erf is the error function.

Specifically, emissions from landfills are estimated by TT using two different methods. The first approach assumes that wood decomposition in landfills occurs under aerobic conditions, with CO_2_ as the only emitted gas (as in Equations 10 and 11). Although this approach may underestimate the warming potential of emissions, it allows for comparison with alternative methods. The second approach is a theoretical first-order decay (FOD) model introduced in the IPCC guidelines (38), which provides yearly estimates of CH_4_ and CO_2_ emissions. The following equations present the generation of CH_4_ and CO_2_ in year *t* from landfilled wood in a specific year *x*.

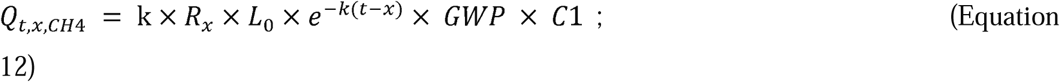

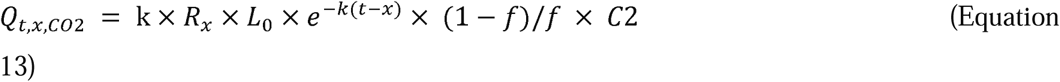

where *Q_t,x,CH4_* and *Q_t,x,C02_* are, respectively, the amounts of methane and carbon dioxide generated in year t (in eq tC) by landfilled wood *R_x_* (t); 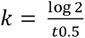 represents the methane generation rate constant (yr^-1^); t0.5 is the half-life period of the degradation process; *L*_0_ is the methane generation potential (tCH_4_ per tonne of landfilled wood); *x* is the year of wood input, and *t* is the current year; GWP = 28 is the global warming potential of CH_4_ (in eq CO_2_), C1 = 12/44 and C2 = 12/16 are the constants used to convert tCO_2_ to tC and tCH_4_ to tC, respectively; and f represents the fraction of C released as CH_4_.

#### 2.5.5. The substitution module

TT simulates the climate change mitigation effect in terms of avoided emissions resulting from the substitution of fossil fuels by bioenergy and energy-intensive materials by wood products. The carbon mitigation potential is closely linked to the specific solution being substituted and is calculated using the displacement factor (DF in tCO_2_-eq m^−3^), a measure of the amount of GHG emissions avoided when wood is used instead of the current solution (39). The computation of the substitution is perfomed separately for material and energy substituiton using the ‘`material_sub()’` and ‘`energy_sub()’` functions, respectively. The substitution is estimated using the following equation:

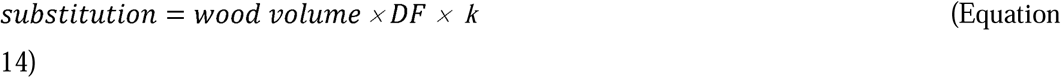

where 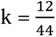 is the constant used to convert tCO_2_ to tC

### 2.6. Case study

#### 2.6.1. Study area

The Bonis experimental watershed located in the mountain area of Sila Greca (39°28′49″ N, 16°32′07″ E; from 975 to 1330 m a.s.l.) in the Calabria region, southern Italy was chosen as the study area in this work (26,40). Almost 93% of its total area is covered by forests, dominated by ~60 years old Laricio pine stands. The stands were planted in 1958 with an average density of 2425 sapling ha^-1^ (41) and underwent a thinning treatment in 1993 with basal area (BA) removal of 25 % (42). The forest was equipped with 14 circular survey plots, each with a radius of 12 meters, for monitoring since 1993, and they were surveyed until late 2016.

**Figure 4:**
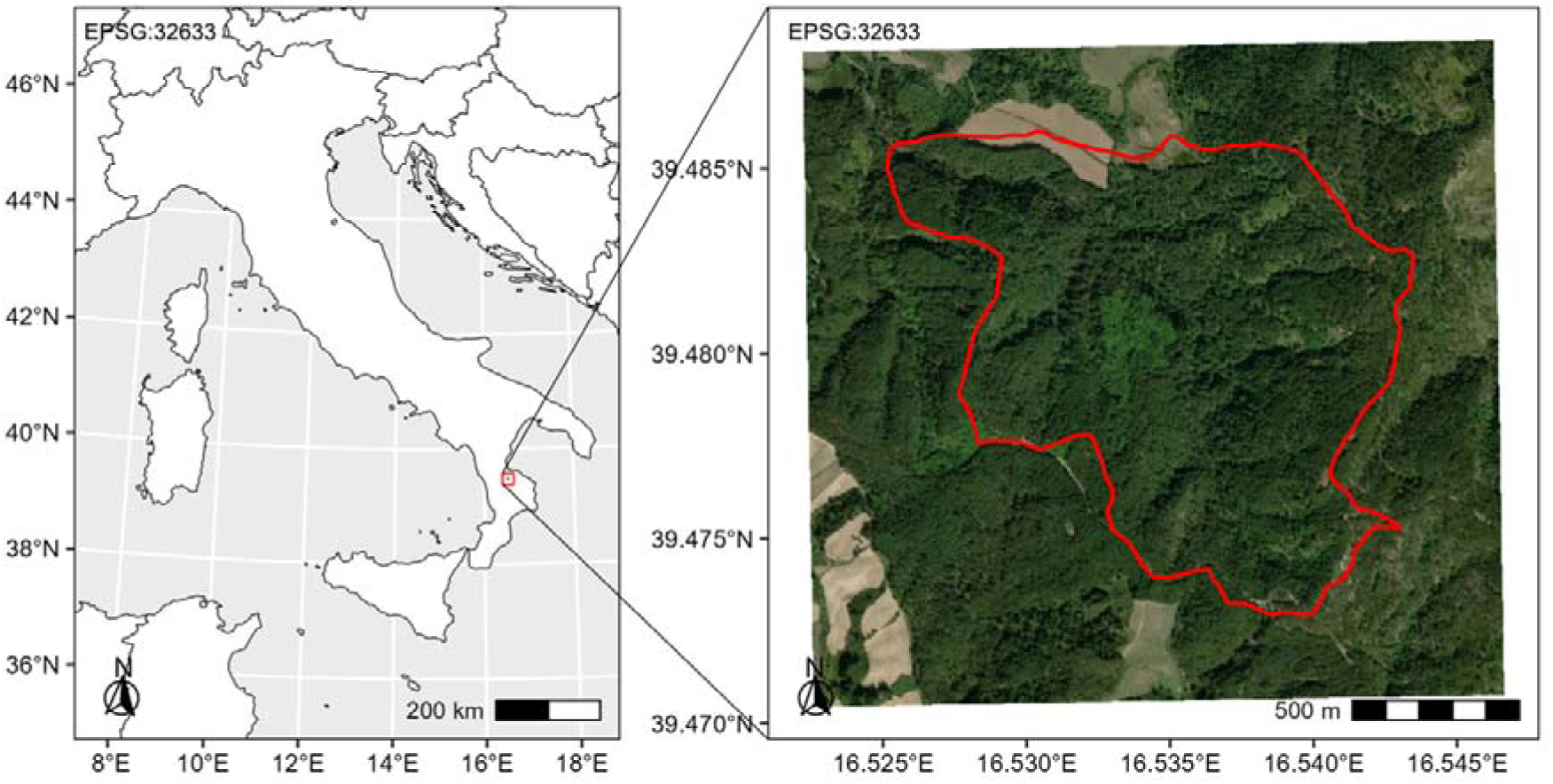
Map of the geographical situation of the Bonis watershed (right sub-figure) inside the Italian territory (left sub-figure).

#### 2.6.2. Scenarios building

For the management scenarios, we tested three options reflecting different goals. All the options were simulated to take place after 2016, which is the last year of plot surveying. The first option simulates light thinning intensity, corresponding to a 28% reduction of Basal Area (BA) at an interval of 15 years, aiming to reproduce silvicultural interventions favoring natural forest dynamics. An additional production-oriented option, known as clearcut, simulates a complete harvest followed by replanting 80 years after the establishment of the plantation. A third option, representing a more sustainable alternative to clearcutting, simulates a shelterwood. This involves two light thinnings (20% reduction of BA) with a 10-year interval, followed by seed-favoring cut after 80 years from the original planting (80% reduction in BA) and removal cut 10 years later.

For the wood-use scenarios, four different options were developed. The ‘Baseline scenario’ (‘Business as Usual’, BAU) kept the recycling rate and products lifespan values constant. In the ‘Longevity scenario’, the lifetime of products was increased by 10 %. In the ‘Reuse scenario’, the recycling rate of products was increased by 10%. In the ‘Sustainability scenario’, both the lifetime of products and the recycling rate were increased by 10% (See **Supplementary 4** for further details).

#### 2.6.3. Modeling framework and the required data

The *TT*, a WPM that tracks carbon in HWPs which was extensively introduced in this paper, was coupled with 3D-CMCC-FEM (‘*Three Dimensional – Coupled Model Carbon Cycle – Forest Ecosystem Module’* v.5.6 BGC (25,26,43–46), a stand-level, biogeochemical, biophysical, process-based model that annually provides data on the forest state (e.g., density, DBH, BA, and total height). The integration of the two models was considered as the modeling framework for achieving the objectives of this study.

The 3D-CMCC-FEM model requires a set of input data for its initialization which includes: (i) model species parameters set which was derived from a recent work that validated the model for Laricio pine stand in the Bonis watershed (40); (ii) daily time series of meteorological fields (e.g. incoming shortwave radiation, maximum and minimum temperature, relative humidity or vapor pressure deficit, precipitation). For the period from 1958 until 1976 climate data was derived using the mountain microclimate simulation model MT-CLIM (47) forced by temperature and precipitation series measured by the nearby Cecita meteorological station (39°23′51″ N, 16°33′24″ E; 1180 m a.s.l.), while for the period from 1976 to 2005, gridded climate time series were used. The latter derived from bias-corrected outputs of the regional climate model COSMO-CLM (48) at around ~8 km horizontal resolution (49,50), and driven by the general circulation model (GCM) CMCC-CM (51) under historical GHG forcing (40). Additionally, measured values of global annual atmospheric CO_2_ concentration were derived from (52) and used for the period from 1958 to 2005. A random sampling of both climate and CO_2_ data within the period between 1990 and 2005 was performed as representative of an additional synthetic period of 90 years assuming unchanging climate and atmospheric CO_2_ conditions, to simulate in total 140 years; (iii) stand initialization data for the year 1958 which included stand density: 2425 saplings ha^−1^, DBH: 1 cm, height: 1.3 m, age: 4 years, elevation: 1131 m a.s.l., soil texture (clay: 20 %; silt: 26 %, sand: 54 %) and depth: 100 cm (41,53,54). (40) provides an extensive description of model validation (before and after thinning) at the Bonis site for both carbon fluxes and stocks such as DBH, stand density and gross primary productivity (GPP).

In addition, *TT* also requires a set of inputs and parameters necessary for its initialization: i- the bucking allocation criteria, developed to segregate solid-wood products based on quality requirements for specific end uses across various wood products, are standardized and can be applied irrespective of log source and sawmill producer (55). Potential wood products from Laricio pine stems were inventoried by consulting five sawmill industry experts, and the amalgamation of all possible products was retained for this study. Furthermore, the bucking criteria used in this study are those commonly found in the literature (56) (Please refer to Table 1 in **Supplementary 2**); ii- the transformation efficiency of each log category, a geometric yield as well as the loss reallocation were defined with the assistance of sawmill industry professionals (please refer to Tables 2 and 4 in **Supplementary 2**),; iii- for the recycling rate we suggest that the recycling rate of waste wood products was constant at 10% during the planning horizon while lifespan of each product was reviewed from published studies (57–63); iv- displacement factors for the substitution were derived from the literature (39,64) (Please refer to Table 5 in **Supplementary 2**); v- TT model dendrometry parameters, including stand structure, crown base height equation, and diameter-to-height parameters, were fitted to the forest data collected from the experimental plots of the Bonis watershed forest, while the taper model and the species density were derived from the literature (65,66).

**Table 2:** Forest management and wood use scenarios.

| Group | Name | Features |
| --- | --- | --- |
| Forest management | Selective thinning | 28% reduction of BA at an interval of 15 years |
|  | Clearcut | Complete harvest followed by replanting at 80 years. |
|  | Shelterwood | Two light thinning (20% reduction of BA) with 10-year interval, followed by seed-favoring after 80 years (80% of BA) and removal cut 10 years later |
| Wood use | BAU | Constant recycling rate and product lifespan |
|  | Longevity | 10% increase in product lifespan |
|  | Reuse | 10% increase in recycling rate |
|  | Sustainability | 10% increase in both product lifespan and recycling rate |

Additionally to the first implementation which assumes static displacement factor values overtime, we dynamically calculated these factors for each product category in this study, following the approach outlined in (21). Specifically, the substitution factor value was assumed to proportionally follow the dynamic of gross anthropogenic CO_2_ emissions. In line with the Paris Agreement’s ambition of reaching net-zero emissions around mid-century, (67) estimated the evolution of these gross anthropogenic CO_2_ emissions. We used this projection to dynamically update the substitution factors throughout the entire 140-year planning horizon.

## 3. Results

### 3.1. Introduction

In this study, we assess the carbon dynamics within the harvested wood product (HWP) pool, including carbon stock changes, emissions, and the effects of material and fossil fuel substitution. We employ the production approach, which tracks the transfer of carbon from forests to HWPs and covers the wood lifecycle from forest harvest to end-of-life. Our analysis focuses exclusively on the C dynamics within the HWP pool, without evaluating changes in forest carbon stocks. This approach is versatile, applicable to various system boundaries, including specific projects and harvest activities of small sizes, including our study scope.

In this simulation, the overall C-balance, calculated as the difference between carbon stored in HWPs, the avoided emissions as effect of the substitution of material and fossil fuel, and carbon emissions from HWPs end-life, was estimated for three different forest management schemes and four wood use scenarios providing insights into the overall carbon balance at each point in time throughout the projection period. These estimations were derived from the modeling exercise, relying on both silvicultural itinerary and product utilization over the use and end-use periods. To analyze the effects of various wood use scenarios on the overall C-balance over time, we compared each scenario with the business-as-usual (BAU) scenario as benchmark. Furthermore, to assess the impact of different forest management scenarios on the overall C-balance over time, we conducted individual comparisons while maintaining the same wood-use scenario each time. Additionally, the net C-balance, calculated as the difference between the carbon stored in HWPs and C-emissions at the end of their life-cycle, was estimated across the three forest management schemes and four wood-use options. This concept is significant because it provides a baseline against which the substitution effect can be measured and can orient comparative studies where substitution is irrelevant. When estimated dynamically, it offers insights into the duration of a positive C-balance, which is crucial for evaluating the effectiveness of HWPs in mitigating climate change. A long-lasting positive C-balance indicates that carbon is sequestered from the atmosphere and stored in HWPs for extended periods, thereby delaying or reducing the impact of C-emissions on global warming. Finally, to assess the impact of considering CH_4_ release from landfill sites and the dynamic updating of substitution factors, we evaluated the effects of incorporating these elements. In this work we use positive sign to represent C-removals while negative one is C-emissions.

### 3.2. Overall carbon balance

The application of wood-use scenarios had distinct effects across different forest management scenarios, yet recurrent patterns can be identified. Among the various forest management scenarios, the sustainability scenario consistently exhibited the lowest C-emissions. To elaborate, over the planning horizon, C-emissions decreased by 69%, 267%, and 40%, respectively, for clearcut, selective thinning, and shelterwood managements compared to the BAU scenario (i.e., –6.1, 1.34, –12.91 tC ha^−1^). Comparable effects were observed with the longevity scenario, resulting in a reduction of C-emissions by 60%, 254%, and 35%, respectively, for clearcut, selective thinning, and shelterwood managements compared to BAU. The reuse scenario induced the smallest decrease in C-emissions, at 11%, 44%, and 4.6%, respectively, for clearcut, selective thinning, and shelterwood managements compared to BAU (see **Table 3**).

**Table 3:** The overall carbon balance, encompassing removals, emissions, and substitution effects from harvested wood products (HWPs) due to wood use and management scenarios, is expressed in tC ha^−1^. Negative sign indicate net C-emissions, whereas positive sign indicates net C-removals.

| Management scenario | Wood use scenario |  |  |  |
| --- | --- | --- | --- | --- |
|  | BAU | Reuse | Longevity | Sustainability |
| Clearcut | -6.1 | -5.43 | -2.46 | -1.88 |
| Selective thinning | 1.34 | 1.93 | 4.74 | 4.92 |
| Shelterwood | -12.91 | -12.32 | -8.39 | -7.76 |

In the context of forest management, selective thinning exhibited pronounced superiority over alternative management approaches in all four wood-use scenarios. Under this management strategy, C-emissions were intensively reduced compared to the BAU scenario for clearcut and shelterwood management, respectively. Meanwhile, clearcut and shelterwood management demonstrated nearly equivalent effects across the three wood-use scenarios (see **Table 3**).

The overall C-balance exhibits diverse patterns based on the applied forest management. Specifically, the results consistently demonstrate a positive balance for the selective thinning management approach throughout the entire planning horizon. However, the clearcut management reached equilibrium at either year 64 or 65 under BAU and the reuse scenarios, or under the longevity and sustainability scenarios. The balance remained negative from that point until a significant harvesting event at year 84. Subsequently, a new equilibrium was achieved at either year 134 or 138 under BAU and reuse scenarios, or under longevity and sustainability scenarios. The overall balance remained negative thereafter. In the case of shelterwood management, the balance consistently remained positive until a first equilibrium was reached at either year 129, 130, or 133 under BAU, or under the reuse scenario, or under the longevity and sustainability scenarios. Afterward, the balance remained negative (See **Figures 5–7**).

**Figure 5:**
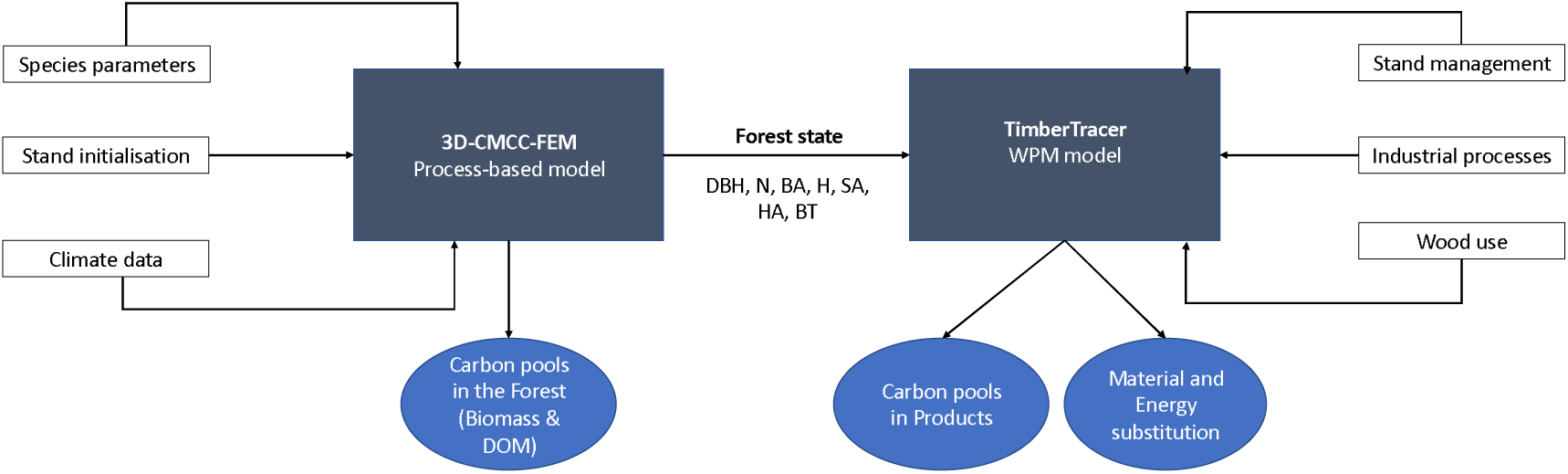
carbon modeling framework coupling a forest process-based model (3D-CMCC-FEM) and TT a harvested-wood product model. DBH is the stand mean diameter at breast height, N is the stand density, BA is the stand basal area, H is the stand mean tree height, SA is the sapwood area of the mean tree, HA is the heartwood area of the mean tree, and BT is the bark thickness of the mean tree.

**Figure 6:**
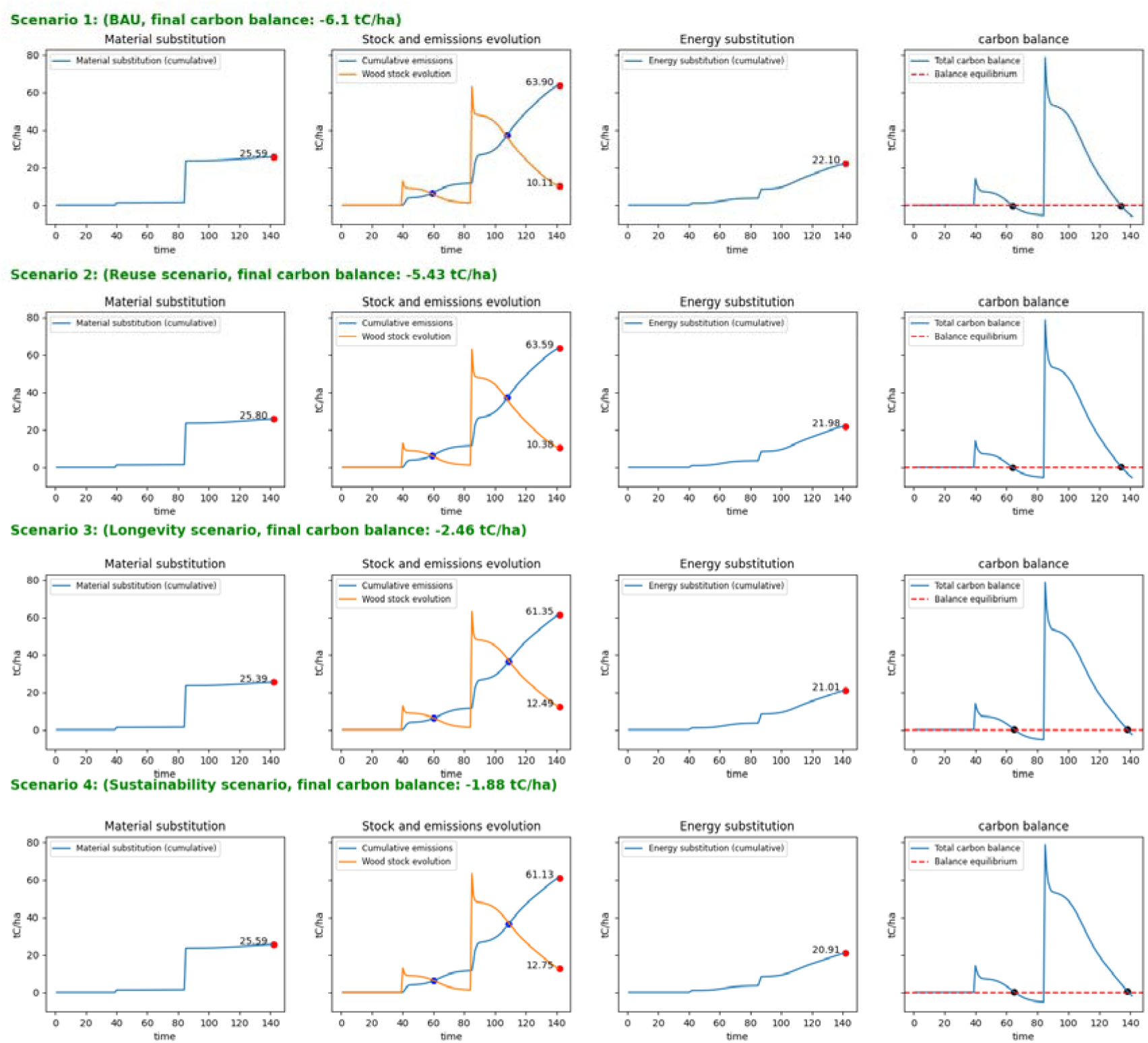
Clearcut management among different wood-use scenarios is demonstrated to showcase variations in Harvested Wood Products (HWPs) C-stock, material and energy substitutions, C-emissions, and the **overall C-balance** (tC ha*^−^*^1^). Blue points represent the equalization between HWPs and emissions, while black points indicate neutrality of the overall C-balance. The red dashed line corresponds to the overall C-balance neutrality. Negative values of the overall C-balance indicate positive emissions, while positive values indicate positive C-storage. The red points refer to the value of each variable at the term of the simulation.

**Figure 7:**
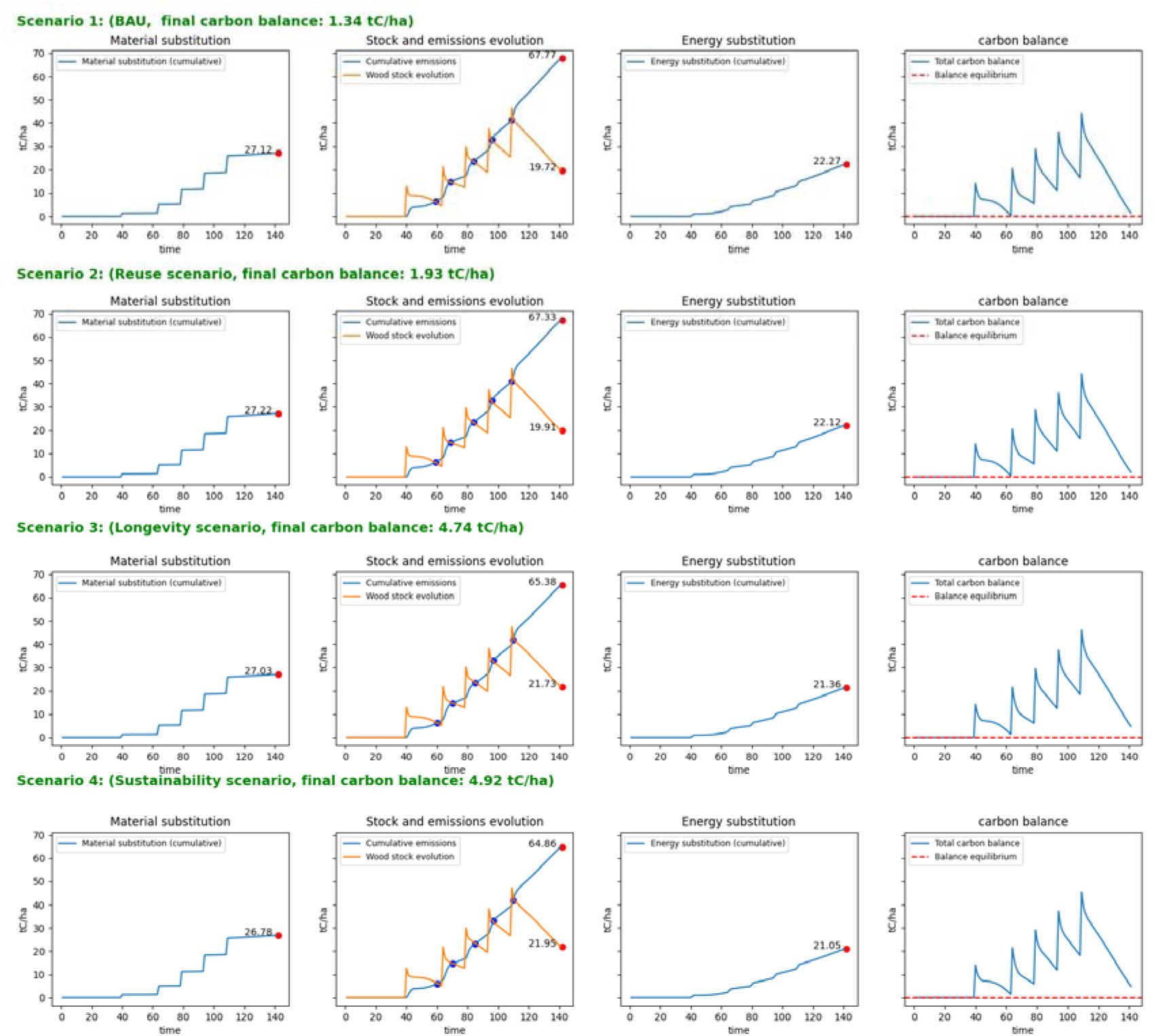
Selective thinning management among different wood-use scenarios is demonstrated to showcase variations in Harvested Wood Products (HWPs) C-stock, material and energy substitutions, C-emissions, and the **overall C-balance** (tC ha*^−^*^1^). Blue points represent the equalization between HWPs and emissions. The red dashed line corresponds to overall C-balance neutrality. Negative values of the overall C-balance indicate positive emissions, while positive values indicate positive C-storage. The red points refer to the value of each variable at the term of the simulation.

**Figure 8:**
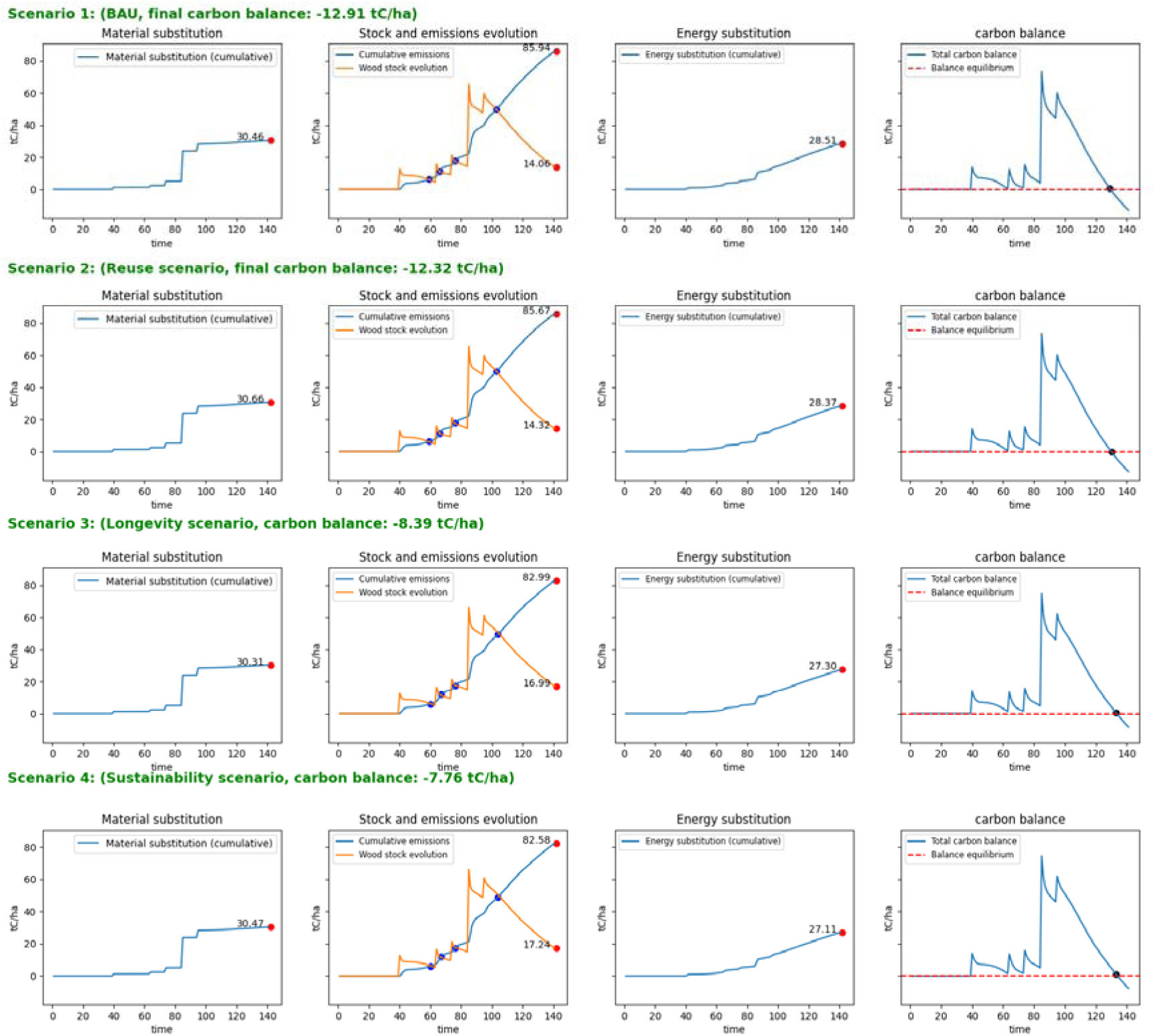
Shelterwood management among different wood-use scenarios is demonstrated to showcase variations in Harvested Wood Products (HWPs) C-stock, material and energy substitutions, C-emissions, and the **overall C-balance** (tC ha*^−^*^1^). Blue points represent the equalization between HWPs and emissions, while black points indicate the neutrality of the overall C-balance. The red dashed line corresponds to the overall C-balance neutrality. Negative values of the overall C-balance indicate positive emissions, while positive values indicate positive C-storage. The red points refer to the value of each variable at the term of the simulation.

### 3.3. Material and energy substitution

Regarding material substitution, we identified a distinct pattern marked by successive phases of a sharp pulse of increase coinciding with thinning or harvesting operations, followed by a positive pseudo plateau. Among the forest management scenarios, the reuse scenario consistently showed the highest material substitution effect averaging 0.62% more than the BAU scenario (i.e., 25.59, 27.12, and 30.46 tC ha^−1^; respectively for clearcut, selective thinning and shelterwood). In contrast, the longevity scenario consistently exhibited the lowest potential (averaging 0.53% less than the BAU) while the sustainability scenario demonstrated a comparable effect to the BAU scenario.

For energy substitution, we observed a pattern characterized by successive phases of a moderately intense pulse of increase coinciding with thinning or harvesting operations, followed by a positive slope. Among the management scenarios, the BAU scenario (i.e., 22.10, 22.27, and 28.51 tC ha^−1^; respectively for clearcut, selective thinning and shelterwood) consistently exhibited the highest energy substitution effect. In contrast, the sustainability consistently showed the lowest energy substitution effect (averaging 5.25% less than BAU), followed by the longevity scenario (averaging 4.41% less than BAU) and finally the reuse scenario (averaging 0.57% less than BAU).

### 3.4. Net carbon balance

Regarding the carbon emissions from firewood and disposal sites, we observed a pattern characterized by a sustained positive slope interrupted by a sharp pulse of increase coinciding with thinning or harvesting operations. Among the management scenarios, the BAU scenario (i.e., –63.90, –67.77, and –85.94 tC ha^−1^; respectively for clearcut, selective thinning and shelterwood) consistently exhibited the highest C-emissions. In contrast, the sustainability scenario consistently showed the lowest level of emissions (averaging 4.17% less than BAU), followed by the longevity scenario (at 3.64% less than BAU), and finally the reuse scenario (at 0.48% less than BAU).

Regarding the carbon stock of HWPs, it assumes different shapes depending on the type of applied forest management. However, a commonly observed pattern is characterized by a sustained negative slope interrupted by a sharp pulse of increase coinciding with thinning or harvesting operations. Among the management scenarios, the BAU scenario consistently exhibited the lowest HWPs C-stock (i.e., 10.11, 19.72, and 14.06 tC ha^−1^ for clearcut, selective thinning, and shelterwood, respectively). In contrast, the sustainability scenario consistently showed the highest level of C-stock in HWPs (averaging 20% more than BAU), followed by the longevity scenario (at 18.2% more than the BAU), and finally, the reuse scenario (at 1.80% more than BAU).

The duration of positive net C-balance differs significantly across various forest management strategies and slightly among different wood-use scenarios (see **Table 4**). In the case of selective thinning, five positive periods, measured in years, can be observed, with their durations decreasing over time: [20, 6, 6, 3, 1] or [21, 7, 7, 4, 2], respectively, for BAU and the reuse scenarios or the longevity and sustainability scenarios. In the case of clearcut management, two positive periods can be observed with their durations slightly increasing over time: [20, 24] or [21, 25], respectively, for BAU and the reuse scenario or the longevity and sustainability scenarios. Regarding the shelterwood management, four positive periods can be observed with their durations decreasing and then increasing overtime: [20, 3, 3, 19] or [21, 3, 3, 20], respectively, for BAU and the reuse scenario or the longevity and sustainably scenarios.

**Table 4:** Duration of positive net C-balance among various forest management and wood-use scenarios (in years).

| Management scenario | Wood use scenario |  |  |  |
| --- | --- | --- | --- | --- |
|  | BAU | Reuse | Longevity | Sustainability |
| Clearcut | 36 | 36 | 41 | 41 |
| Selective thinning | 44 | 44 | 46 | 46 |
| shelterwood | 45 | 45 | 47 | 47 |

### 3.5. Implementation of dynamic substitution and methane release from landfill

Accounting for dynamic substitution and methane release from landfills had a significant impact on the overall C-balance of harvested wood products (HWPs) across all forest management and wood-use scenarios, leading to a substantial decrease in this balance, ranging from 5- to 39-fold compared to the original projections (see **Figure 9**). Moreover, under this new implementation, all combinations of forest management and wood-use scenarios resulted in a negative overall C-balance, including selective thinning, which had presented a positive overall C-balance under the original implementation (see **Figure 9**).

**Figure 9:**
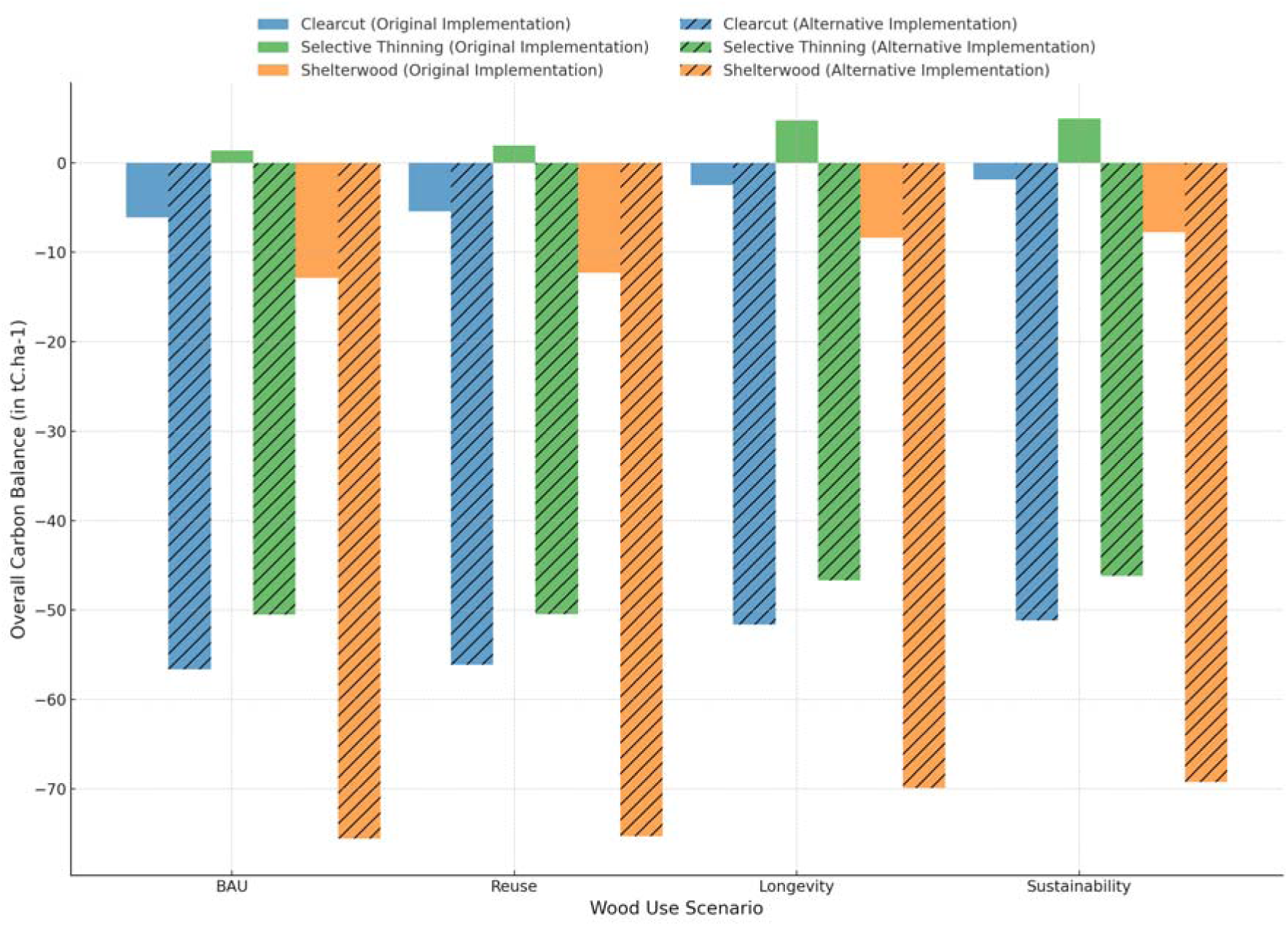
Comparison of the overall C-balance across forest management and wood use scenarios using two implementations: original (no methane (CH_4_) release, static substitution) and alternative (methane (CH_4_) release, dynamic substitution). Negative values indicate net C-emissions.

In contrast, this new implementation had no effect on the ranking of different forest management scenarios, as selective thinning consistently demonstrated superiority over alternative approaches across all wood-use scenarios, followed by clearcut management and, finally, shelterwood management (see **Figure 9**). Similarly, the implementation did not alter the ranking of overall wood-use scenarios, with the reuse scenario showing the minimal climate mitigation potential across all management scenarios, while both the longevity and sustainability scenarios made comparable and higher contributions to C-emissions reduction compared to the BAU scenario (see **Figure 9**).

In the following subsections (**4.5.1** and **4.5.2**), the results related to the impacts of dynamic substitution and methane release will be presented individually, respectively.

#### 3.5.1. Effect of dynamic substitution

The implementation of dynamic updating of displacement factors, proportional to the reduction in global emissions consistent with the Paris Agreement, led to a decrease in displacement factors after reaching their maximum (see Figure A.1). This, in turn, resulted in a significant reduction in the substitution effect across all forest management and wood-use scenarios compared to the initial projections that utilized static displacement factors. By the planning horizon (year 140), cumulative material substitution in the dynamic scenario was reduced by 95% to 96% compared to the static substitution scenario, and cumulative energy substitution in the dynamic scenario was reduced by 95% compared to the static substitution scenario (see **Figure 10**).

**Figure 10:**
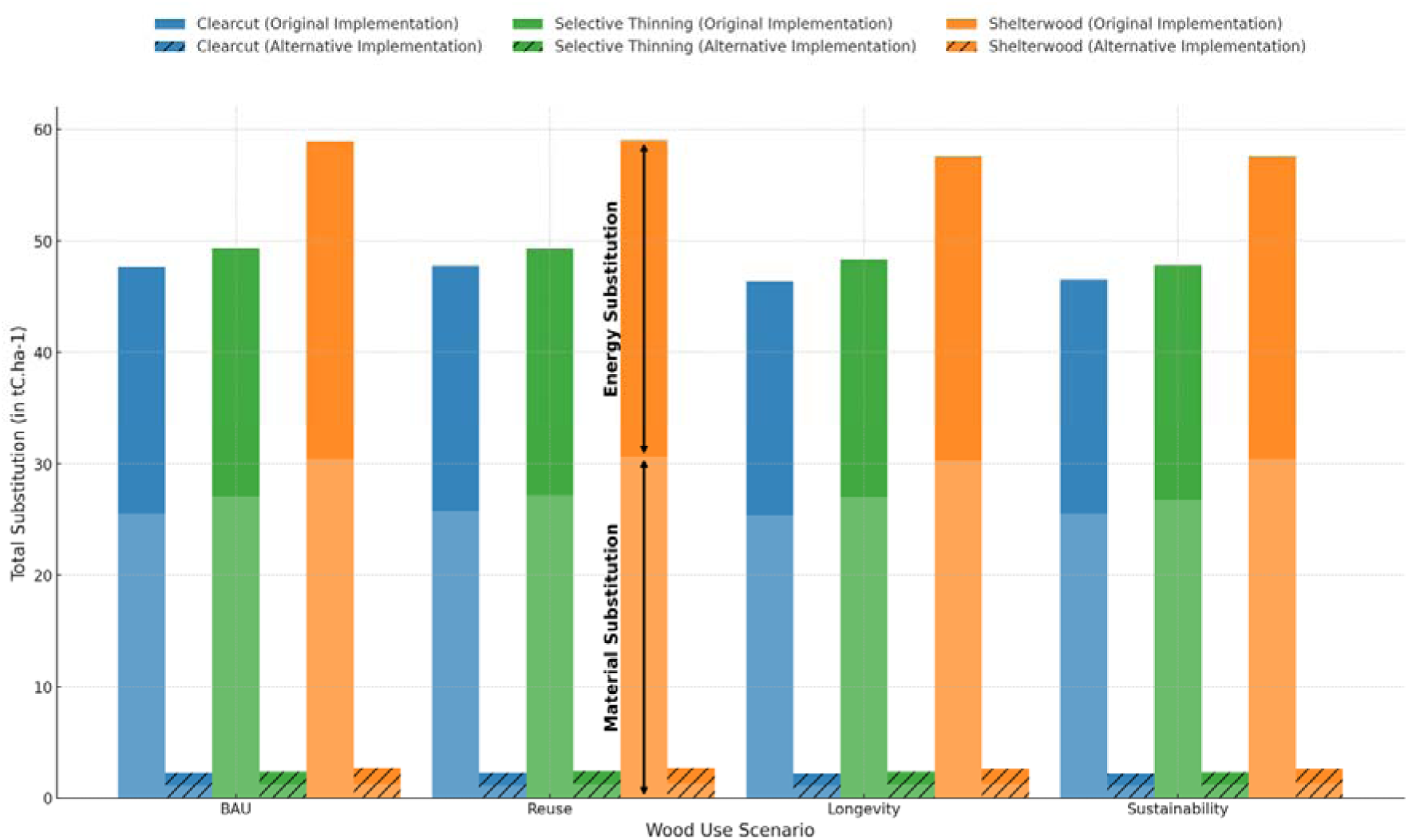
Comparison of the cumulative sum of material and energy substitutions resulting from the use of a particular forest management and wood-use scenario using two implementations: the original (static substitution) and the alternative (dynamic substitution) (in tC ha^−1^). The shaded portions of the bars represent the relative contribution of material substitution, while the solid portions represent the relative contribution of energy substitution.

#### 3.6.1. Effect of methane release from landfills

The implementation of methane (CH_4_) release from landfills under anaerobic conditions, following the IPCC’s theoretical first-order methodology, resulted in a considerable increase in cumulative emissions across all forest management and wood-use scenarios compared to the initial simulations, which only considered wood decomposition in landfills under aerobic conditions. Consequently, by the planning horizon (i.e., year 140), cumulative emissions in the new implementation increased by 9.4% to 11% compared to the original implementation (see **Figure 11**).

**Figure 11:**
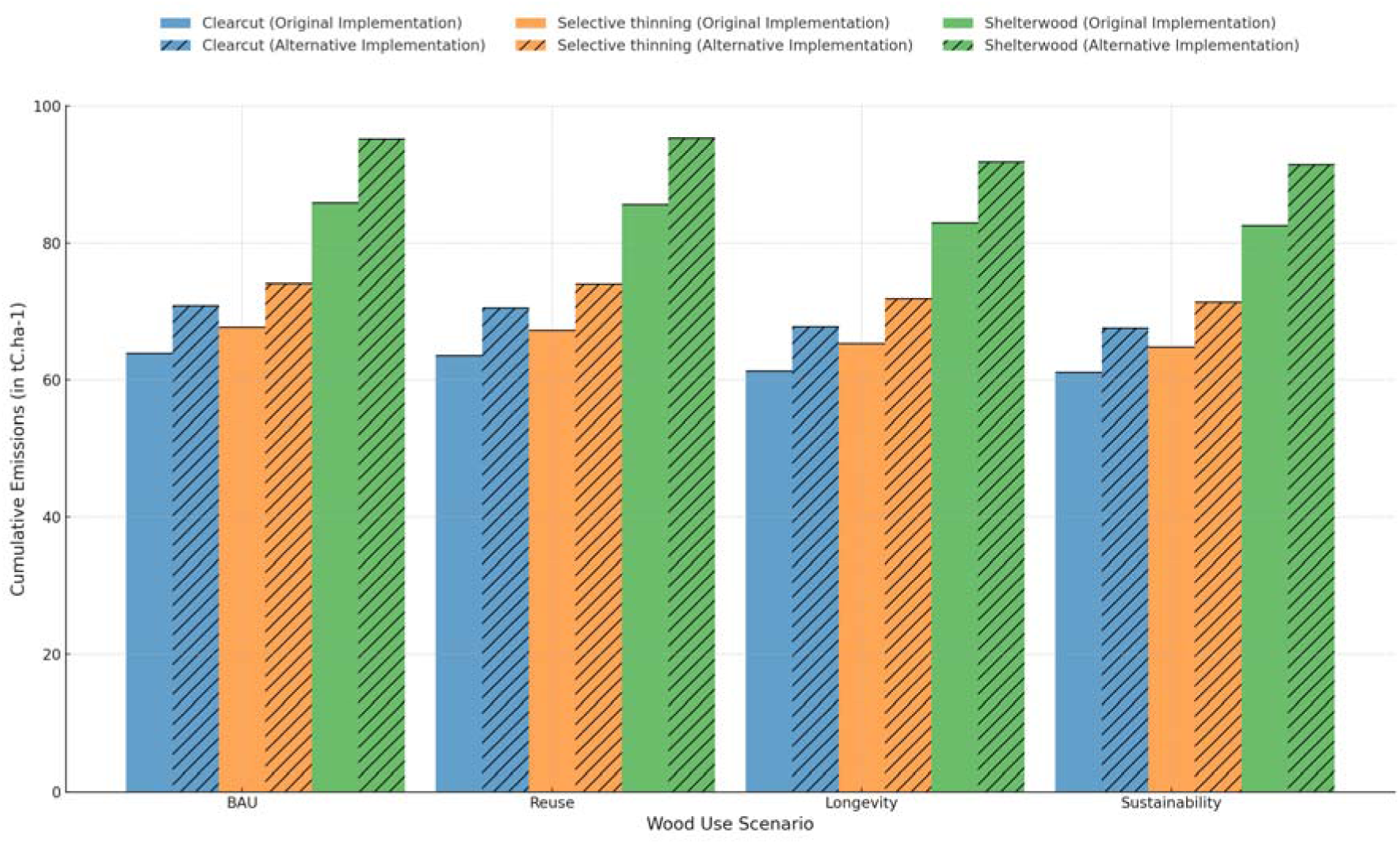
Comparison of the cumulative emissions resulting from the use of various forest management and wood-use scenarios under two implementations: the original (no methane (CH_4_) release) and the alternative (with methane (CH_4_) release) (in tC ha^−1^).

## 4. Discussions

In this study, we presented the underlying theory and principles of TT, a dynamic model of the carbon balance in HWPs. TT has been coupled with 3D-CMCC-FEM with the aim to investigate the effect of different forest management and wood use options on the overall C-balance of HWPs on a real case. Assuming flexibility in both wood utilization and forest management practices, this study case demonstrates that the overall C-balance can be increased by giving preference to multiple light, non-distant cuttings over a few distant intensive cuttings and promoting wood use for material purposes, including the increase of recycling and products lifetime. These findings align closely with the principles of continuous cover forestry and the circular economy approach, emphasizing the role of sustainable resource management in climate mitigation. Additionally, to evaluate the impact of methane emissions from landfills under anaerobic conditions and to account for the dynamic nature of displacement factors, we implemented an alternative model configuration. This approach revealed that considering CH_4_ emissions significantly increased total emissions, while applying dynamic displacement factors substantially reduced the substitution effect. However, these findings did not alter the relative ranking of forest management and wood-use strategies in terms of their climate change mitigation potential.

Recent researches increasingly questions the static nature of substitution factors, advocating for a dynamic approach that accounts for technological advancements, improved resource efficiency, enhanced recycling processes, and better end-of-life management of products (e.g., (18,20)). The substantial reduction in substitution effects observed in this study highlights the sensitivity of carbon accounting to displacement factor assumptions, reinforcing the need for adaptive modeling approaches. Consistent with our results, (21) demonstrated that applying dynamic displacement factors leads to a projected 33% reduction in the mitigation effect of wood product substitution in Europe by 2030, and a 96% reduction by 2100, compared to predictions using fixed factors. Similarly, (18) conducted simulations showing that product substitution would saturate if displacement factors reached zero within 100 years of the simulation’s start, resulting in an 89% reduction in cumulative substitution effects within 300 years. These findings emphasize the need for continuous reassessment of displacement factors to ensure an accurate representation of substitution benefits.

Methane emissions from landfills are reported under the waste sector according to IPCC guidelines for reporting emissions and sinks, which is why the methane effect is often excluded from the calculation of HWP contributions by most WPMs. However, accounting for methane emissions is crucial for developing practices that enhance the contribution of HWPs, especially given methane’s high global warming potential. Even a minor underestimation of CH_4_ can lead to a significant and erroneous overestimation of the potential of various mitigation projects. In this study, including CH_4_ release from landfills due to wood decomposition under anaerobic conditions led to a substantial increase in emissions (in CO2 equivalents), ranging from 9.4% to 11% compared to the original implementation where methane emissions were not considered. Consistent with our results, (68) demonstrated that accounting for the global warming potential of methane emissions reduced the estimated contribution of U.S. forests and HWPs to GHG mitigation for the year 2005 by more than half (in C-equivalent). However, it should be noted that the inclusion of landfill gas management infrastructure or the adoption of strategies, such as the ongoing EU initiative aiming to limit municipal waste landfilling to below 10% by 2035 as part of the broader goal of achieving climate neutrality by 2050, may substantially reduce CH_4_ emissions. As such, WPMs should account for these considerations (69).

The outcomes of this study are significantly enhanced by the inclusion of the bucking allocation module in the TT model, which considers the dimensions of logs for the allocation of wood to HWPs. This is crucial because various products have distinct use and post-use properties, leading to diverse temporal dynamics, and a pre-established allocation may not accurately reflect reality. The role of the bucking allocation was evident across all management options. Each successive intervention, characterized by a higher mean DBH of the harvested stems than the precedent due to the stand’s growth dynamic, resulted in the newly HWPs stock exhibiting a slower retirement dynamic than the precedent due to the higher portion of long-lived products in the most recent one. For instance, in the case of clearcut management, two interventions were made at the years 39 and 84, yielding retirement rates of ~2% per year and 1.5% per year, respectively. The analysis of the bucking allocation in both the initial and subsequent harvests, reveals that, from the first harvest, the production was limited to short- and medium-lived HWPs such as paper and particle, while novel categories of long-lived products were introduced in the second harvest, exemplified by furniture and sawing, which justify the decrease in the retirement rate of the HWPs stock (see **Table A.1**).

These multiple findings align with prior studies in this area. In a theoretical exercise evaluating the mitigation potential of wood product use in the European forest sector, (17) demonstrated that increasing each component—whether recycling rate or lifespan— individually by approximately 20% could result in an 8.9% increase in C-removal by 2030 by reference to the 2017 BAU scenario. Furthermore, the study states that a simultaneous 20% increase in both average product lifespan and recycling rate could yield a 17.3% increase in carbon removal by 2030. Another recent study by (70) explored the C-storage potential of HWPs in four EU countries, projecting outcomes from 2020 to 2050 across six alternative scenarios. The findings suggest that prioritizing wood-use for material purposes, while maintaining a constant harvest, yields the highest mitigation benefits in the short to medium term. Moreover, in a continental study (EU-28) focusing on assessing the consequences of implementing policy choices on GHG emissions and removals, it was revealed that the adoption of the cascading scenario of HWPs led to a slight increase in the net balance between emissions and removals from/by HWPs. The balance was simulated to rise from approximately 34 Mt CO_2_-eq in the base period around 2010 to just under 40 Mt CO_2_-eq in 2030, as documented by (71). In another study aiming to investigate the potential of cascading use of woody biomass in the EU, it was found that GHG emissions could be reduced by 35 MtCO_2_-eq year^−1^ as a result of implementing the maximum technical potential to increase recycling of waste wood and paper flows (72).

From another perspective, the significance of management has been underscored in numerous studies. In (73), the application of five silvicultural itineraries, derived from translating five alternative management objectives for an even-aged forest, resulted in significantly different outcomes regarding the carbon storage by HWPs. In another study, (74) concluded that the method of harvesting is crucial showing further that a regular removal of timber from forest in a way that maintains a continuous canopy is likely to give substantially higher sustained yields and amount of carbon sequestration than periodical clear-felling because of the higher photosynthetic area. The same study suggests that if the objective was to maximize timber volume yield, the optimal management system would be the regular thinning of forest. A physiological explanation of this could be that the continuous canopy cover with a moderately high leaf area index (~ 4 m^2^ m^−2^) provides high light interception and, at the same time, net primary production (75). Regular thinning ensures that the forest has lower biomass than an undisturbed forest, and it is continuously growing, resulting in lower maintenance respiration (4). In a different context, (76) demonstrated that selection harvesting was the preferred method compared to clearcutting when the goal was to maximize total carbon stock in the forest landscape and wood products generated from harvesting over an 80-year planning horizon. This preference was justified by the fact that selection harvesting, in contrast to clearcutting, offers the advantage of maintaining the forest close to its maximum biological productivity because of maintaining the better and more productive trees. Additionally, it provides a consistent and sustainable yield of desirable wood products at set intervals. In contrast, clearcutting involves harvesting stands when their average DBH reaches 10 cm (the common merchantable dimension), and their yields exceed 50 m^3^ ha^−1^. This practice is more likely to favor the production of pulpwood due to the smaller size of the harvested trees, which implies a faster retirement of the derived products and thus of the HWPs stock.

## Limitations and perspectives

Considering the model’s sensitivity to inputs and the dependence of results on the approach used for calculating HWPs stock, emissions, and substitution, it is crucial to account for uncertainties associated with these elements (77). The TT model tracks wood throughout its entire lifecycle, from harvesting to disposal sites, thereby encompassing major processes in between. This modeling principle is considered advanced due to its capacity to accurately trace carbon over the lifetime of wood products, providing precise results. However, implementing this principle at the national level poses challenges due to the large number and diversity of HWPs, as well as the substantial amount of the required local data (15). As the amount of data increases, the level of compounded uncertainty proportionally rises. A subsequent phase would involve scrutinizing the sensitivity of various model outputs to the input data. Additionnaly, uncertainty arising from the coupling of 3D-CMCC-FEM and TT must be carefully considered. Given the feedforward nature of this coupling, the individual uncertainties of each model reflect their potential impact on the integrated framework outputs leading, in the best scenario, to compensation errors or, in the worst, to their sum. Model uncertainties generally stem from three primary sources: parameter values, which represent process importance; model structure, which defines how these processes are represented; and the initial state, which sets the starting conditions for the simulation (78,79). These uncertainties are particularly important as the outputs of 3D-CMCC-FEM serve as inputs for TT. The dynamics of the 3D-CMCC-FEM model have been extensively studied and are known to vary across forest development stages and tree species. Early growth is primarily driven by structural parameters such as plant architecture, light interception, and LAI, which indirectly influence photosynthesis. As the forest matures, long-term processes such as carbon allocation, autotrophic respiration, and biomass accumulation become dominant, influencing overall biomass and net primary productivity (NPP) (45). This time-dependent uncertainty in model parameters highlights the need for fine-tuning the 3D-CMCC-FEM to account for different time scales. Moreover, the uncertainty of 3D-CMCC has been evaluated based on hypotheses representing key physiological processes. For example, in (80), two plant respiration theories—photosynthesis-dependent and biomass-dependent—were assessed against observational data. This study revealed that respiration is not fully governed by either theory, prompting revisions to how 3D-CMCC estimates the respiration.

From another perspective, the grading of logs utilizing the bucking allocation module, as exemplified by the implementation in TT, encounters obstacles due to various factors presumed to impact wood quality — such as strength, knottiness, appearance, stiffness, hardness, and durability. It is almost impossible, using exclusively models, to account for all these factors. As an illustration, the implemented module does not account for the presence of residual branches below the crown base and external defects. Furthermore, it is conceivable that variations in knot distribution exist across different management scenarios. Owing to the omission of this parameter in our model, we assert that the disparities in the proportions of HWP classes could be somewhat undervalued. This conjecture remains subject to empirical verification, yet the current capabilities of TT do not permit the necessary resolution for such details. Therefore, future refinements of the model are imperative to enable a more precise evaluation of these nuances. Furthermore, as demonstrated in (16), the choice of one distribution function over another to simulate product removal from use can significantly influence the resulting carbon stock estimates. The TT model currently utilizes the normal distribution as its default set up; however, this selection should be reassessed in future analyses.

Given that the maximum climate benefit varies over time for different forest managements (81,82), we raise questions about the relevance of the fixed planning horizon of 140 years in this study and its capacity to appropriately address the underlying research hypothesis. In this context, we may explore in the future the potential of alternative management strategies, such as clearcut and shelterwood, to be optimal for different planning horizons. Furthermore, we may question the assumption that simulations conducted over multiple rotations can effectively control for instantaneous response.

Additionally, the substitution effect is subject to considerable uncertainty due to various methodological and market-related factors. (18) identifies key assumptions that can lead to an overestimation of substitution benefits, including the use of static displacement factors, the assumption that no leakage or cross-sectoral material replacement occurs, and the notion that building longevity and substitution longevity are independent. Similarly, (20) highlight uncertainties related to market behavior, particularly whether an increase in wood production translates into higher HWP consumption and whether wood-based materials consistently remain economically competitive. (83) emphasize the role of market demand, consumer behavior, and policy frameworks in determining whether substitution benefits can be realized. (84) considers the socio-economic factors influencing substitution, noting that cultural preferences and regulatory barriers may limit large-scale adoption of wood products. Finally, (85) highlight that economic cycles can significantly affect HWP carbon storage, as market downturns or reduced construction activity may slow wood consumption, temporarily reducing substitution benefits or even turning HWPs into a net carbon source. These studies collectively highlight the need for dynamic modeling approaches that account for technological progress, evolving market conditions, and real-world constraints rather than relying on simplified, static assumptions.

Moreover, it is crucial to note that carbon neutrality does not necessarily imply climate neutrality. When wood is burned or decays, emissions spend some time in the atmosphere before being sequestered, contributing to climate change in the meantime. In other words, the timing of emissions and sinks has an impact on the overall cumulative climatic impact (86–88). Among the factors affecting this timing are the speed of biomass regrowth (rotation period) and the storage of biomass products (e.g., building, furniture, and paper). Since these factors are closely linked to forest management practices, the assessment of these practices should not only consider the carbon balance but also examine the timing of carbon input and output flows.

Another question concerns the methodological scheme used in this study. Evaluating the potential of various forest management and wood-use options in mitigating climate change necessitates a systemic perspective that considers the diverse pools of the forest sector, including biomass, soil, and products (89,90). The current study exclusively focuses on the wood products pool, which may not be directly relevant (or being the only one) for guiding the decision-making of forest managers. This is particularly important considering that the forest ecosystems constitutes the major contribution of the forest sector in terms of climate change mitigation. To exemplify this recent assertion, (9) calculated the emissions and removals linked to HWPs for the historical period (1992-2012) and future scenarios until 2030 in the EU (excluding Malta and Cyprus). They utilized FAOSTAT data on forest product production (see https://www.fao.org/faostat/en/#data/FO). The findings of this research indicate that the average historical sink of HWPs from 2000 to 2012 accounts for 10% of the sink contained in the forest pools. This underscores the importance of including non-HWP carbon pools in decision-making processes as well.

Finally, the results of this use case suggest that shelterwood management is the optimal choice for optimizing the overall carbon balance of HWPs within the defined planning horizon. The rationale behind this preference lies partially in the late thinning operation characteristic of this management scenario, allowing for the C-stock and substitution to counterbalance total emissions effectively. Notably, this silvicultural practice of partial cutting aligns with natural disturbance–based silviculture, as it mimics natural dynamics by anticipating the imminent mortality of a portion of mature trees (91). Moreover, considering the ongoing rapid changes in many ecosystems and their interaction with natural disturbances, which are expected to be significant and less predictable in the future (92), multi-aged forest management systems offer a promising approach to enhance resistance and resilience, which is attributed to the presence of multiple age classes, providing more potential pathways for post-disturbance management and recovery (93). Within this context, a suggestion arises to guide future studies in accounting for disturbance risks. Neglecting to incorporate such considerations could result in an overestimation of the climate mitigation efficacy associated with various forest management alternatives. The emerging results show that the failure to account for methane release from landfills can lead to an overestimation of the mitigation potential of different options. Efforts to mitigate these emissions could focus on diverting wood products away from landfills by increasing recycling and directing them to incineration facilities, which not only reduce emissions but also capture energy. The results also indicate that considering static substitution factors may overestimate the substitution effect resulting from the use of HWPs over fossil energy or energy-intensive materials. However, we believe that *ceteris paribus*, substitution could play a central supporting role in the absence of a strong mitigation policy.

## 5. Conclusions

This research emphasizes the pivotal role of harvested wood products (HWPs) in climate change mitigation, as recognized in various Nationally Determined Contributions (NDCs) under the Paris Agreement. Our novel framework, ‘TimberTracer’, coupled with the 3D-CMCC-FEM forest growth model, enables dynamic assessment of forest management practices and wood-use scenarios, revealing selective thinning as the optimal forest management practice. Additionally, increasing recycling rates and extending product lifespans prove effective in enhancing C-balance. Results show also that failure to account for methane release from landfill and dynamic substitution of HWPs can lead to an overestimation of the mitigation potential of different options. These insights offer valuable guidance for advancing knowledge and optimizing forest management decisions and advancing climate change mitigation efforts.

## Data availability

The data resulting from this study is available in both Supplementary Files 1 and 4. Additionally, the output simulation is accessible via the following Zenodo link: https://zenodo.org/records/10692189

## Competing interests

Authors have no competing interests as defined by BMC, or other interests that might be perceived to influence the results and/or discussion reported in this paper.

## Code availability

TimberTracer is a Python based model for all operating systems (Windows, Linux, and MacOS). It is free and open source (version 1.0.0 with GPL-3 license, requiring Python >= 3x). We openly share our model on GitHub for collaborative research, fostering a community-driven approach to innovation. A tutorial on Google Colab empowers users to harness TimberTracer’s capabilities, customize analyses, and integrate it seamlessly into projects. Detailed instructions are available on GitHub (https://github.com/issamyax/TimberTracer), emphasizing TimberTracer’s primary objective: providing valuable insights into carbon storage, substitution, and emissions progression over time. We encourage users to report any issues and/or desired extensions on our active issues page https://github.com/issamyax/TimberTracer/issues. The 3D-CMCC-FEM model code version 5.6 is publicly available under the GNU General Public Licence v3.0 (GPL) and can be found on the Github platform at https://github.com/Forest-Modelling-Lab/3D-CMCC-FEM.

## Funding

I.B and R.V were supported by the National Research Centre for Agricultural Technologies — AGRITECH — PNRR (Italian National Plan of Recovery and Resilience), CN00000022 in WP 4.3 Task 3 Risk management strategies and policies in the context of climate change. A.C and D.D acknowledge the project funded under the National Recovery and Resilience Plan (NRRP), Mission 4 Component 2 Investment 1.4 - Call for tender No. 3138 of December 16, 2021, rectified by Decree n.3175 of December 18, 2021 of Italian Ministry of University and Research funded by the European Union – NextGenerationEU under award Number: Project code CN_00000033, Concession Decree No. 1034 of June 17, 2022 adopted by the Italian Ministry of University and Research, CUP B83C22002930006, Project title “National Biodiversity Future Centre - NBFC

## Authors’ contribution

I.B: Conceptualization, Data curation, Formal analysis, Investigation, Methodology, Resources, Software, Validation, Visualization, Writing – original draft. A.C, S.L, and D.D: Resources, Writing-review & editing. Farid Nakhle: Software, Writing-review & editing. R.T: Resources, Writing-review & editing. MV.C and M.S: Resources, Writing-review & editing. R.V: Conceptualization, Supervision, Formal analysis, Methodology, Writing-review & editing, Funding acquisition.

## Acknowledgements

I.B. would like to thank the University of Tuscia and the Euro-Mediterranean Center on Climate Change, both based in Italy, for their support in conducting his Ph.D. thesis, supervised by Professor Riccardo Valentini, of which this work is a part.

## Consent for publication

I hereby grant permission for the publication of my work. I confirm that I have the authority to grant this permission.

## Ethics approval and consent to participate

I consent and confirm that ethics approval has been obtained.

## Appendix A

**Table A.1:**
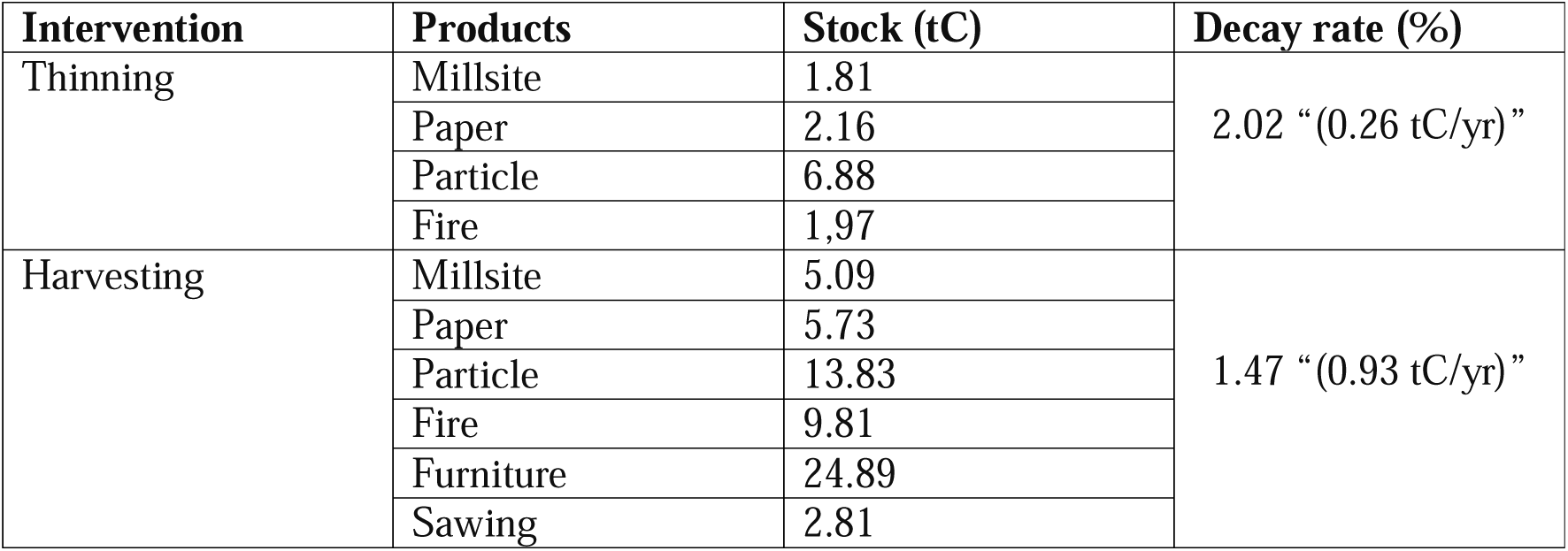
Clearcut management interventions and the products derived from the thinned wood.

| Intervention | Products | Stock (tC) | Decay rate (%) |
| --- | --- | --- | --- |
| Thinning | Millsite | 1.81 | 2.02 “(0.26 tC/yr)” |
|  | Paper | 2.16 |  |
|  | Particle | 6.88 |  |
|  | Fire | 1.97 |  |
| Harvesting | Millsite | 5.09 | 1.47 “(0.93 tC/yr)” |
|  | Paper | 5.73 |  |
|  | Particle | 13.83 |  |
|  | Fire | 9.81 |  |
|  | Furniture | 24.89 |  |
|  | Sawing | 2.81 |  |

**Figure A.1:**
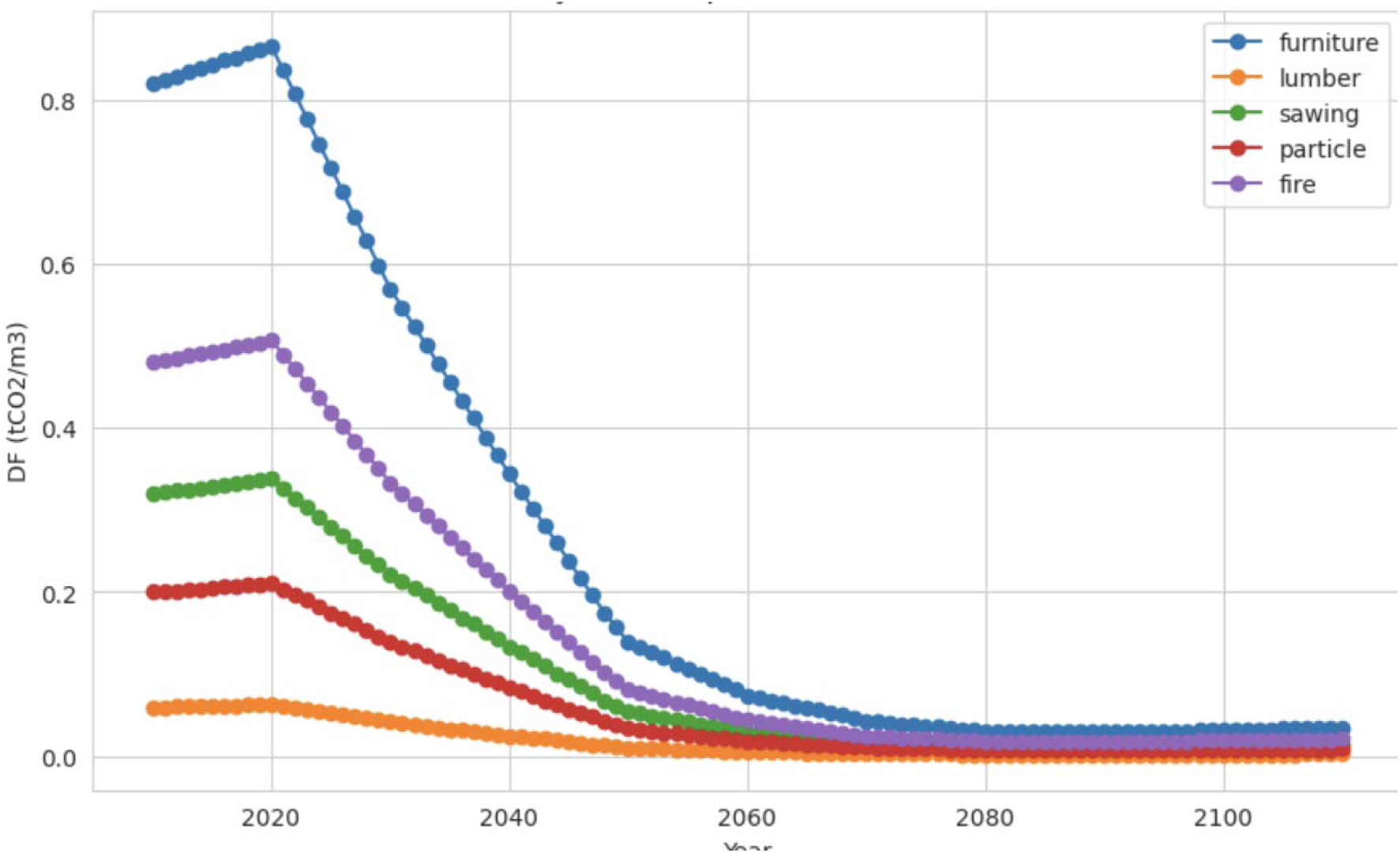
Dynamic evolution of substitution factors of different HWPs proportionally to the evolution of global GHG emissions following the Paris Agreement pathways.

## Notes

### Competing Interest Statement

The authors have declared no competing interest.

### Summary of Updates

Modifications and improvements mainly concerned the introduction and discussion.

