## Supplementary material for "*TimberTracer*: A Comprehensive Framework for the Evaluation of Carbon Sequestration by Forest Management and Substitution of Harvested Wood Products": S1 (from another version of this preprint: version 3, 2025-01-15)

**S1: *TimberTracer* v.1.0.0 - Theoretical framework and concepts**

In supplementary material S1, we introduce the theoretical foundations of *TimberTracer*. Given the modular architecture of the model, we conduct a detailed analysis, examining each module individually. Each of the five modules is accompanied by a graphical representation illustrating the procedural sequence (Figures 1,4,7,8,10), followed by an in-depth exploration of the associated theory and functions.

**1. The dendrometer routine**

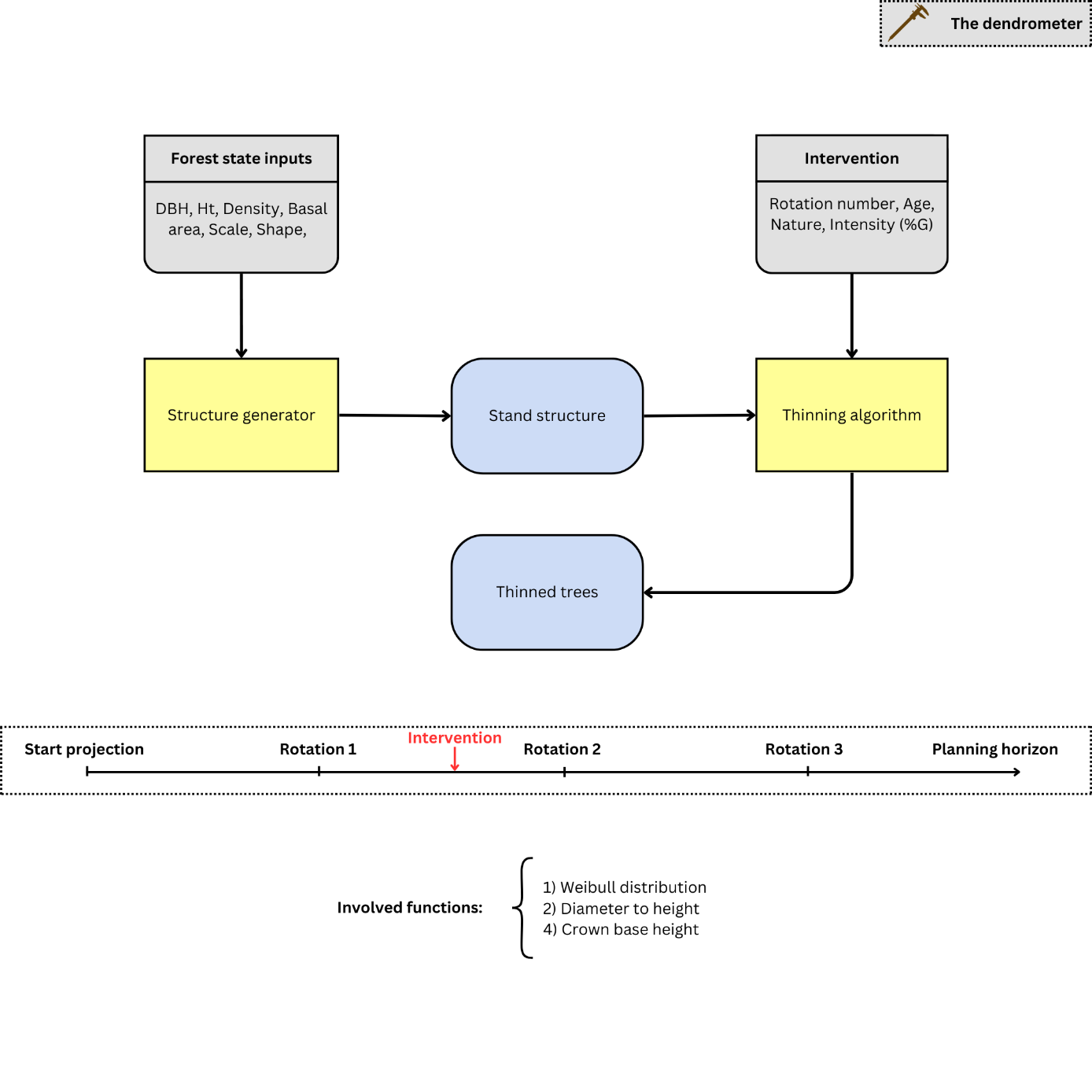

Figure 1: Graphical representation of the dendrometer routine

**1.1. Structure Generator**

The transition from stand state descriptors to tree state descriptors would be feasible if the stand structure is well understood. The latter is typically likened to a statistical distribution. There are numerous mathematical formulations for statistical distributions in forestry, with one of the most commonly employed being the Weibull distribution. The probability density function of a Weibull random variable denoted as $X \sim Weibull(sh, sc, loc)$ is:

$f\left( x, sh, sc, loc \right)=\frac{sh}{sc}{\times\left( \frac{x-loc}{sc} \right)}^{sh-1}\times e^{-\left( \frac{x-loc}{sc} \right)^{sh}}$	(Equation1)

Where f is the probability density function, x is the random variable, sh represents the shape, sc denotes the scale, and loc indicates the location.

If the location is not provided, but the shape and scale are (2-parameter Weibull distribution), then the location will be predicted using the following equation:

$E\left( x \right)=loc+sc \times\Gamma(1+\frac{1}{sh})$	(Equation2)

$loc=E\left( x \right)- sc \times\Gamma(1+\frac{1}{sh})$	(Equation3)

Where E(x) is the mathematical expectancy of x and $\Gamma$ is the gamma function.

After fitting the parameters of the Weibull distribution supposed to represent the structure of the forest stand at different stages of development, the next step is to compute the number of trees by each diameter class. For this specific purpose, the cumulative distribution function (CDF) of the Weibull function is applied to the concerned diameter class following the equation:

$N\left( class \right)= \left[ e^{{-\left( \frac{lower bound-loc}{sc} \right)}^{sh}} - e^{{-\left( \frac{upper bound-loc}{sc} \right)}^{sh}} \right]\times N$	(Equation4)

For an illustrative demonstration see the figure 2 below:

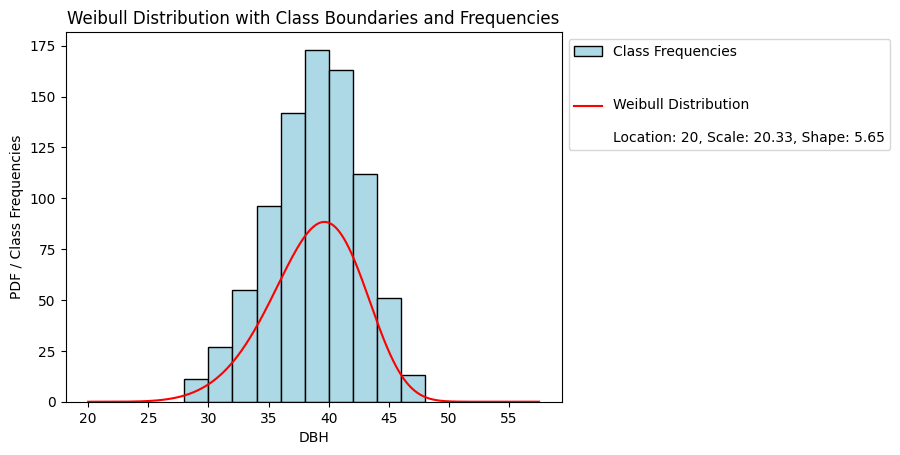

Figure 2: Graphical representation of a 3-parameter Weibull distribution of stand DBH

**1.2. Thinning Algorithm**

A thinning operation involves removing stems to benefit a tree or a group of trees deemed essential to ensure optimal growth conditions. It focuses on enhancing wood quality and stand stability. The thinning cut can be characterized by three elements: i- **the type of cut** (from the above, from the bottom, or neutral), which denotes the distribution of removed stems among various diameter categories of standing trees within the stand ii- **the thinning intensity** which expresses the magnitude of the extraction conducted within the stand. It can be quantified by both the number of stems harvested and the implementation rate relative to the before-thinning stand population iii- **the period**, commonly referred to as rotation, is the time interval between two successive thinning cuts within the stand. It varies depending on the tree species, age, and site conditions.

Bio-sociological tree status plays an important role in some thinning concepts, for example, for the selection of thinning trees in thinning from above and below. A simplified classification groups trees into four categories: **1-** dominant trees **2-** co-dominant trees **3-** intermediate trees and **4-** overshadow trees. This classification consists of the definition of the position of each tree in terms of its height by reference to the other trees of the stand. As part of the process, the dominant height of the plot (**h_95%_**) as well as the height to the base of the crown (**hc**) are calculated. The classification of the trees into a scale is performed following the rules presented in the Table 1 and illustrated in Figure 3.

Table 1: Bio-sociological status criteria based on the tree height.

| **Criteria** | **Bio-sociological status** |
| --- | --- |
| $hi \geq h_{95\%}$ | Dominant tree (1) |
| $\frac{h_{95\%}+hc}{2}\leq hi< h_{95\%}$ | Co-dominant tree (2) |
| $hc\leq hi< \frac{h_{95\%}+hc}{2}$ | Intermediate tree (3) |
| $hi<hc$ | Overshadow tree (4) |

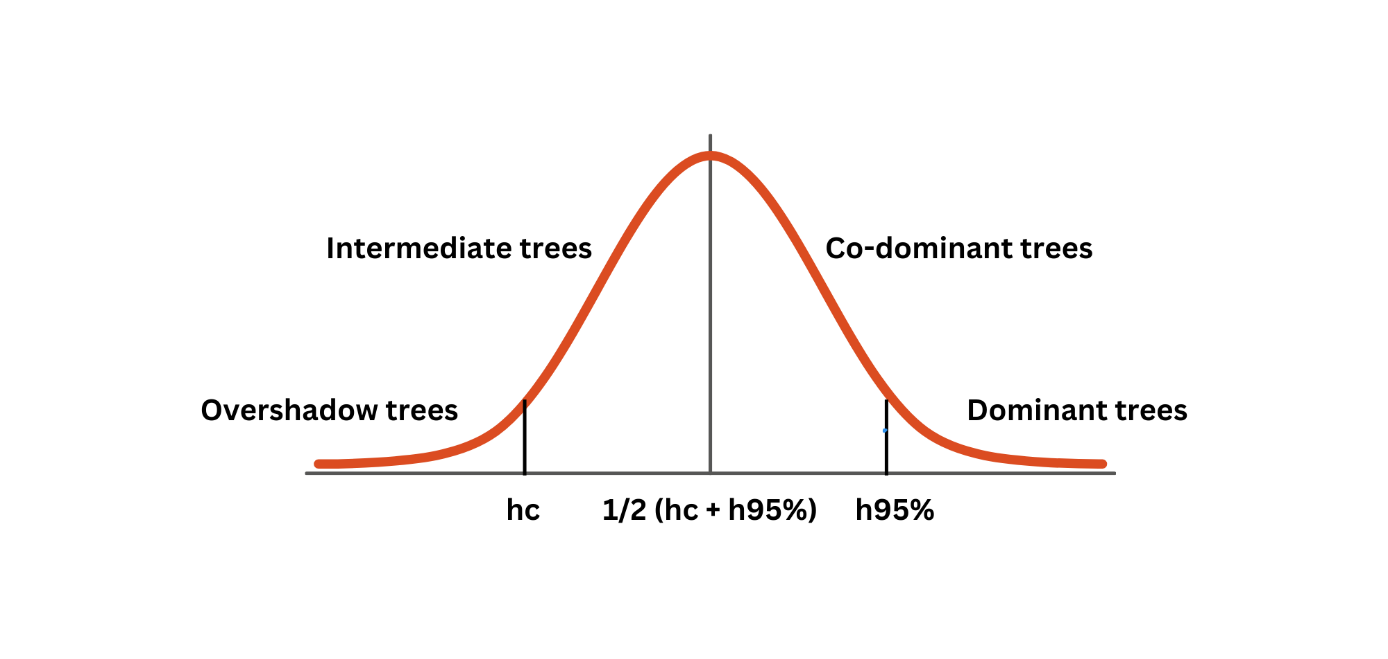

Figure 3: Graphical representation of the bio-sociological tree status and the height thresholds

The Table 2 in the following presents the structure of the results generated using the structure generator.

Table 2: Example of the data frame of the whole trees of the stand

| **Tree_ID** | **DBH (cm)** | **ht (m)** | **SC** | **G (m^2^)** |
| --- | --- | --- | --- | --- |
| 1 | d_1_ | h_1_ | 4 | g_1_ |
| 2 | d_1_ | h_1_ | 4 | g_1_ |
| N | d_k_ | h_k_ | 1 | g_k_ |

**Thinning types:**

**Thinning from below (4+3) => 2**

The process consists of removing trees belonging to the dominated bio-sociological sub-groups. The process of removing is parallel in 4+3 (Overshadow and Intermediate). If the removal amount is not reached by 4+3, the process continues sequentially in sub-group 2 until reaching the required number of trees satisfying the initial condition.

**Thinning from above (1+2) => 3**

The process consists of removing trees belonging to the dominating bio-sociological sub-groups. The process of removing is parallel in 1+2 (Dominant and Co-dominant). If the removal amount is not reached by 1+2, the process continues sequentially in sub-group 3 until reaching the required number of trees satisfying the initial condition.

**Neutral thinning (1+2+3+4)**

The process consists of removing trees regardless of the bio-sociological sub-group to which they belong. The process is parallel in 1+2+3+4 and continues until satisfying the required removal amount.

The Table 3 in the following presents the structure of the results generated using the thinning algorithm.

Table 3: Example of the data frame of the thinned trees

| **Tree_ID** | **DBH (cm)** | **ht (m)** | **SC** | **G (m^2^)** |
| --- | --- | --- | --- | --- |
| 7 | d_1_ | h_1_ | 4 | g_1_ |
| 16 | d_1_ | h_1_ | 4 | g_1_ |
| 79 | d_3_ | h_3_ | 4 | g_3_ |
| N | d_k_ | h_k_ | 1 | g_k_ |
|  |  |  |  | **%G ± tolerance** |

**2. The bucking allocation routine**

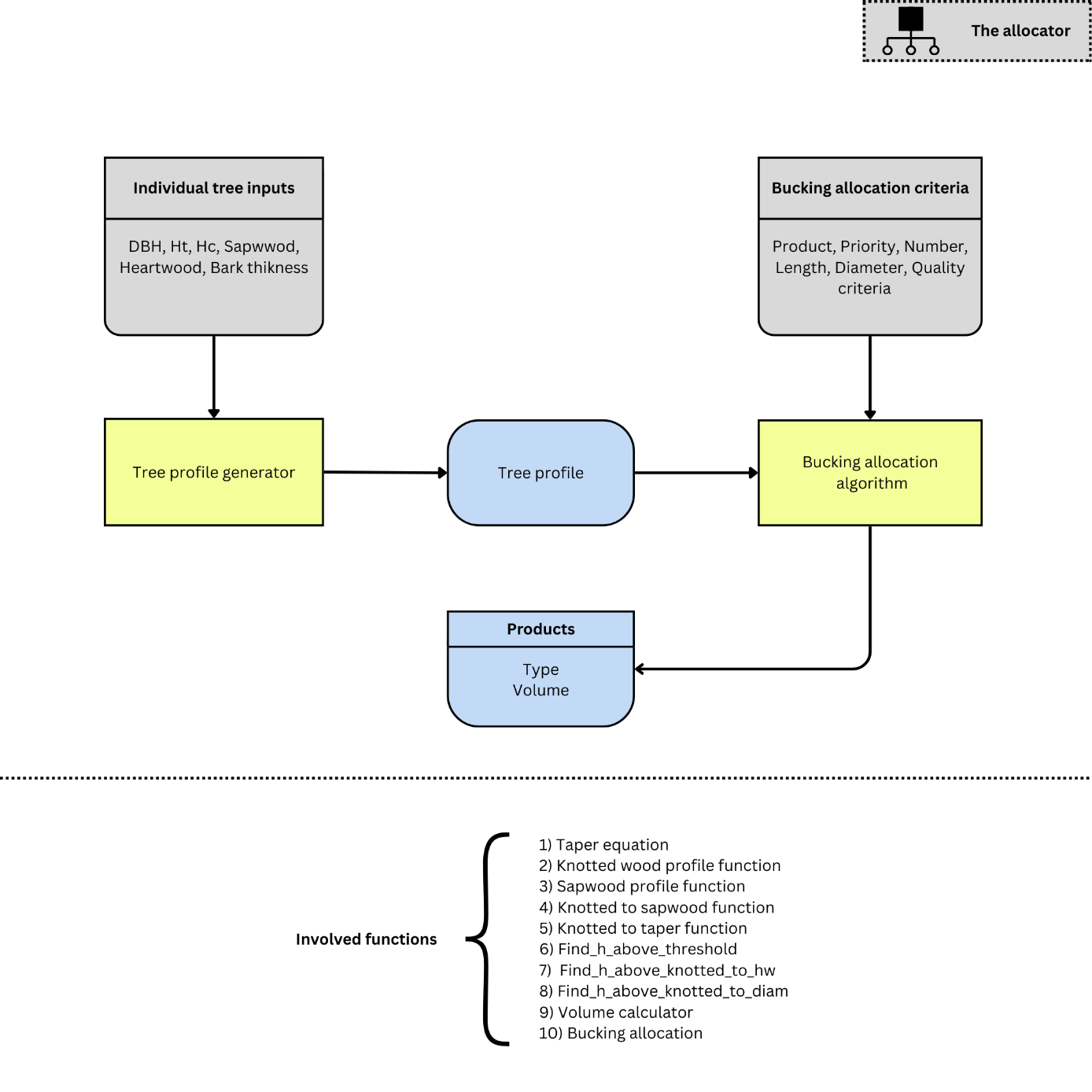

Figure 4: Graphical representation of the bucking allocation module

The bucking allocation refers to the allocation of stems to logs destinated to produce specific harvested products. Determinant factors include species, stem diameter, and wood quality. Practically, this operation is performed following two sequential steps. The first step consists of dressing the **stem profile** including the components for tree quality assessment, while the second step consists of combining the stem profile outputs with the **bucking allocation** criteria to disaggregate the tree into different products.

**2.1. Stem profile generator**

To achieve precise stem profile dressing, this model uses a comprehensive set of equations, which will be thoroughly described in this section.

**taper profile**

Refers to the degree to which a tree’s stem bole decreases in diameter as a function of height above ground. Taper is often represented by mathematical functions fitted to empirical data, called taper equations. One such function, attributed to Max and Burkhart (1976) is:

$y\left( z \right)=b_{1}z+b_{2}z+ b_{3}\left( z-a_{1} \right)^{2}I_{1}+ b_{4}\left( z-a_{2} \right)^{2}I_{2}$	(Equation 5)

**crown base height**

Is an allometric function that derives the crown base height knowing the total stem height. One such function is the 1st-degree polynomial function expressed as follows:

$h_{c}={ah_{t}}^{c}$	(Equation 6)

**total height to DBH**

Is an allometric function that derives the DBH of a tree knowing its total height. One such function is the 1^st^-degree polynomial function expressed as follows:

$dbh=alpha{{\times h}_{t}}^{beta}$	(Equation 7)

**knotted wood profile**

In tree analysis, a critical operation involves dressing the crown base profile to distinguish knotted from intact wood. This function collaborates with three other key functions: i) the taper equation, ii) the crown base height function, and iii) the DBH-to-height relation, collectively facilitating a comprehensive analysis of tree structure.

This system equation consists simply of the reconstruction of the historical crown base height limits and considers that the right side of the curve is the knotted wood and the left side of the curve is the intact wood.

$z_{c}=1-a({h_{c}}^{c-1})$	(Equation 8)

$d_{c}=\sqrt{y(z_{c})}\times alpha \times{{[(\frac{h_{c}}{a})}^{\frac{1}{c}}]}^{beta}$	(Equation 9)

**sapwood_heartwood**

This function computes the sapwood width of a tree by leveraging the sapwood and heartwood areas provided by the growth model.

sap_width = $\sqrt{\frac{4}{\Pi}\left( sap_{area}+heart_{area} \right)}$ - $\sqrt{\frac{4}{\Pi}\left( heart_{area} \right)}$ (Equation 10)

**heartwood profile (Sapwood profile in the model)**

This function computes the heartwood profile under the simplifying assumption that sapwood and bark widths are constants along the stem.

e = bark + sap_width ; bark refers to the bark thickness

diff = dx - e

d_heart = diff; diff > 0

d_heart = 0; diff ≤ 0

The ensemble of the functions defined above is graphically presented in Figure 5.

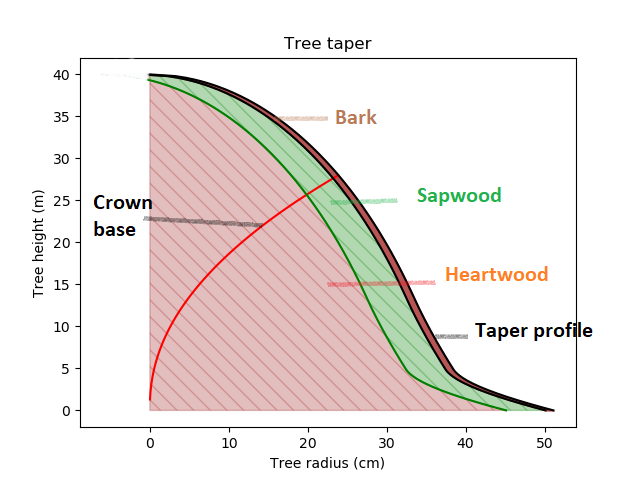

Figure 5 : Graphical representation of the tree longitudinal profile

**2.2. Bucking allocation algorithm**

Logs are disaggregated into various products based on specific criteria, typically defined by wood industry professionals. These criteria encompass both dimensions and quality considerations. Log dimensions are characterized by the length and small end diameter, while log quality is assessed through ratios of knotted wood to heartwood and knotted wood to small end diameter. The Table1 in Supplementary 2 provides an example of cutting rules applicable to Pinus nigra species.

**find_h_above_treshold**

This function implements the bisection method to determine the height level that meets a specified diameter condition (e.g., ØSE = 7 cm or ØSE = 25 cm).

**find_h_above_knotted_to_hw**

Also utilizing the bisection method, this function calculates the height level that satisfies the ratio of knotted wood to heartwood condition (e.g., (Ø_KW_ / Ø_HW_)^2^ ≤ 13%).

**find_h_above_knotted_to_diam**

Like the previous functions, this one uses the bisection method to compute the height level that satisfies the ratio of knotted wood to taper condition (e.g., Ø_KW_/ Ø_SE_ ≤ 30%).

**volume_calculator**

This function calculates the stem volume between two defined heights by integrating the previously defined taper equation.

**bucking allocation**

This method combines the functions described above and user-defined cutting rules to segment the stem into various products.

Figure 6 in the following section graphically demonstrates the disaggregation of the stem into different logs destined to produce specific HWPs, following the bucking criteria outlined in Table 1 of Supplementary 1. Meanwhile, Table 4 presents the structure of the results generated using the bucking allocation algorithm.

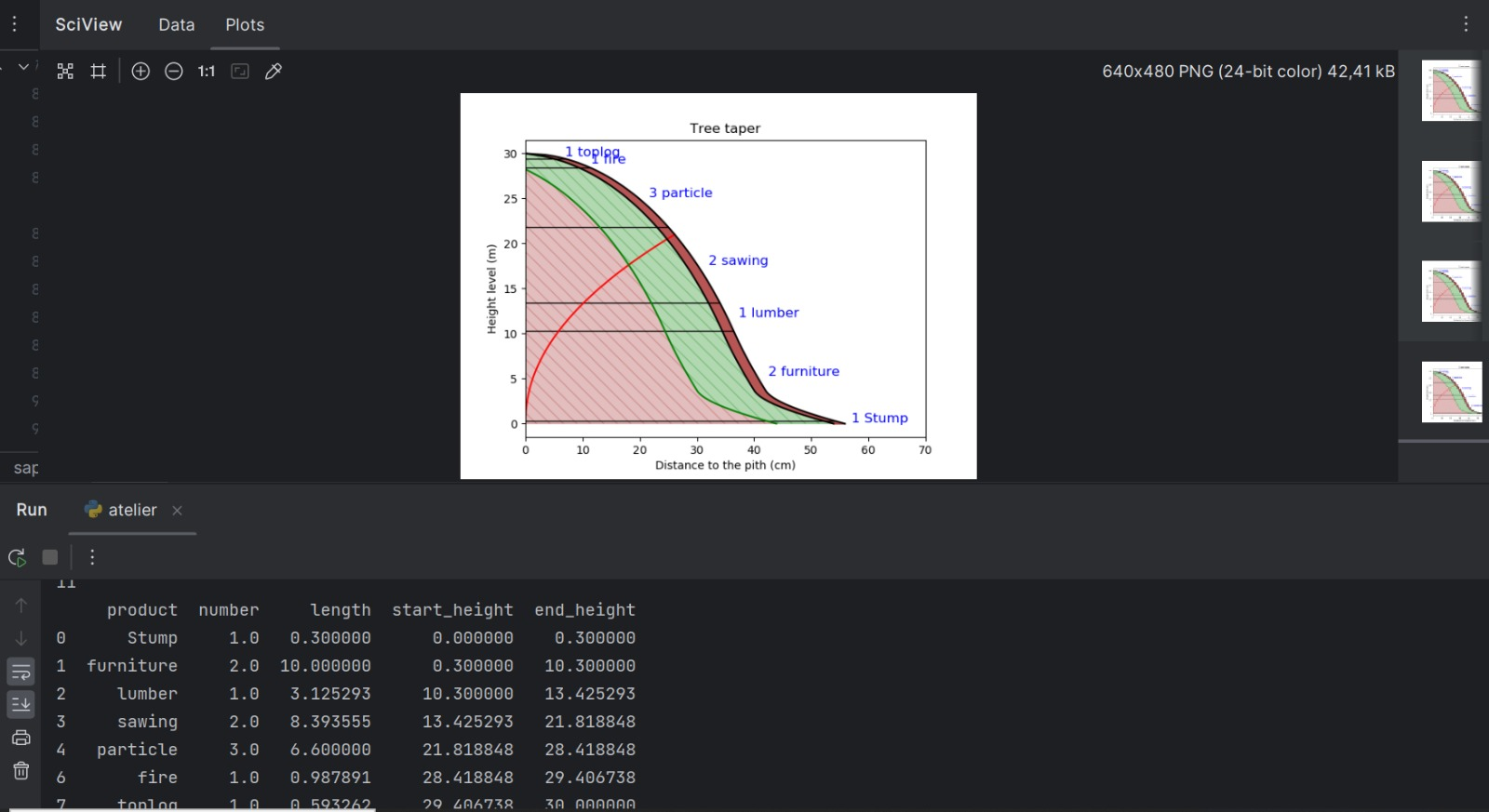

Figure 6: Graphical representation of the stem aggregation into logs allocated to the production of various HWPs.

Table 4: An example of result structure from the bucking allocation algorithm applied a tree of 30 m height.

| **Product** | **Number** | **Start height (m)** | **End height (m)** | **Length (m)** | **Volume (m^3^)** |
| --- | --- | --- | --- | --- | --- |
| Stump | 1 | 0 | 0.30 | 0.30 | - |
| Furniture | 2 | 0.30 | 10.30 | 10 | - |
| Lumber | 1 | 10.30 | 13.42 | 3.12 | - |
| Sawing | 2 | 13.42 | 21.01 | 8.39 | - |
| TopLog | 1 | 29.40 | 30 | 0.60 | - |

**3. The processor routine**

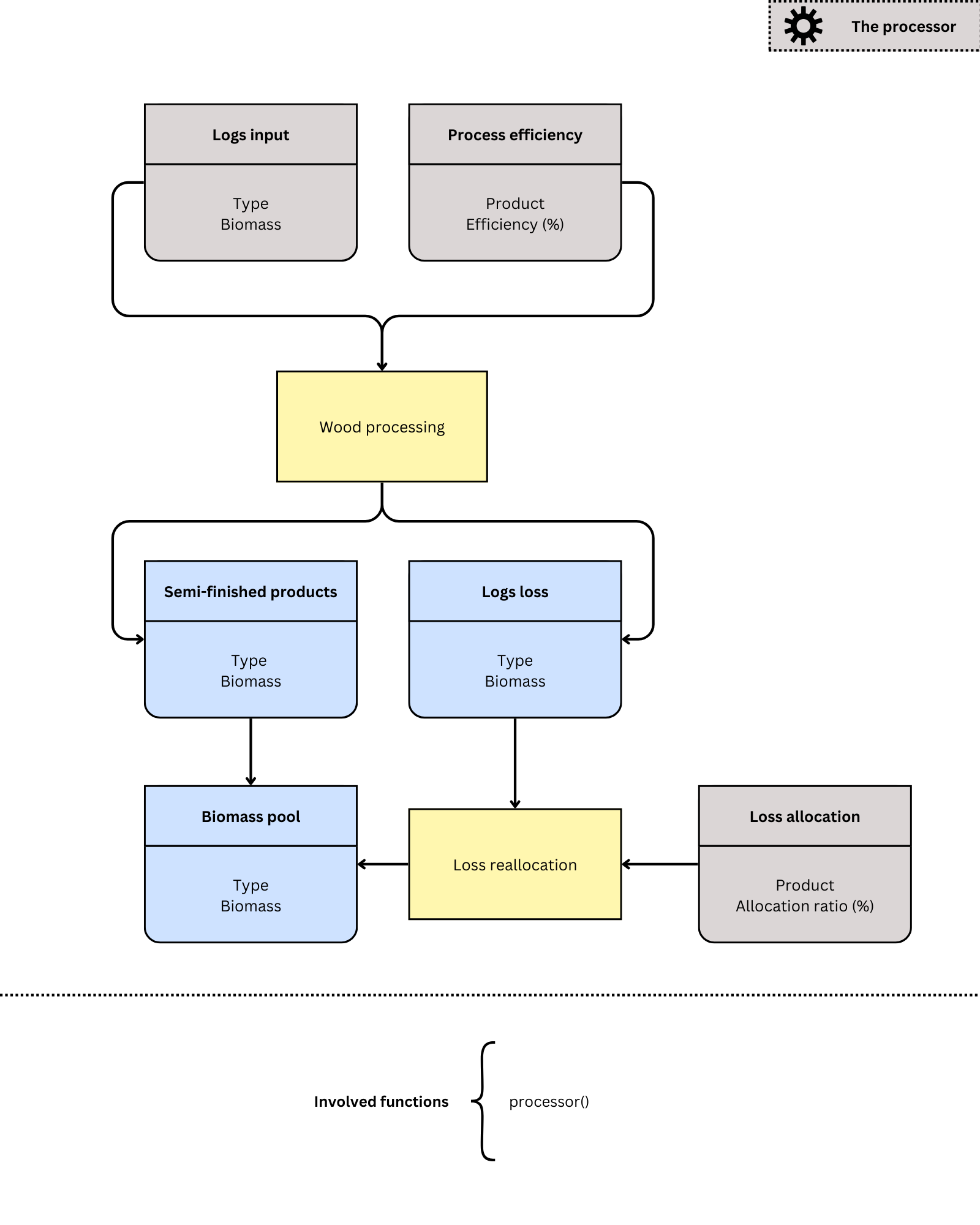

Figure 7: Graphical representation of the processor routine

The processor routine considers the efficiency of producing wood products from logs. Each harvested wood product is linked to a streamlined log transformation process, defining its material efficiency. Process losses are then either redistributed to products of lower grade or discarded at the millsite, with proportions specified by the user. Examples of process efficiency and loss allocation for *Pinus nigra* are detailed in Tables 2 and 3 in Supplementary 2, respectively. In the *TimberTracer* model, this process is implemented in the **processor()** function. The Table 5 below presents the structure of results generated using the processor function.

Table 5: An example of the result structure rendered by the processor function.

| **Product** | **volume** | **post_process_volume_** |
| --- | --- | --- |
| Furniture | - | - |
| Lumber | - | - |
| Firewood | - | - |

**4. Stock and flow dynamic routine**

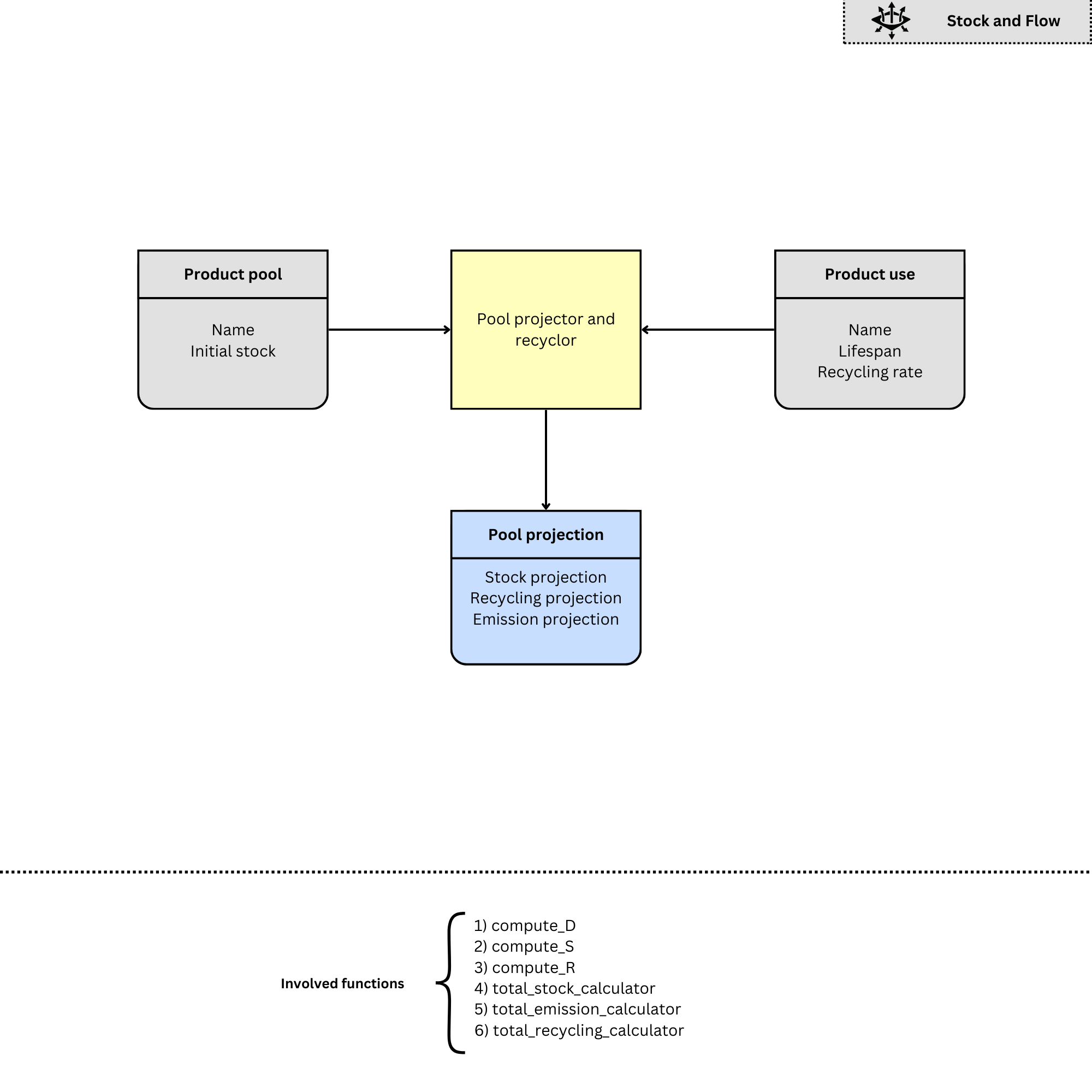

Figure 8: Graphical representation of the stock and flow routine

This module monitors the carbon fate within the HWPs pool, considering the inflow from new wood provisioning and the recycling of products at the end of their use, and the outflow from emissions from firewood and HWPs at disposal sites.

The product removal rate is determined using the cumulative distribution function (CDF) of a normal distribution, which is defined by its mean and standard deviation. The mean represents the average lifespan of a product, while the standard deviation reflects the change in the dynamic decay, typically expressed as a fraction of the lifespan (e.g., a fraction of 1/3 as used in Brunet-Navarro et al., 2017). In the following, the CDF formulation is presented.

$CDF\left( x; lifespan, lifespan \times\mathrm{fraction} \right)=\frac{1}{2} [1+\mathrm{erf} \left( \frac{x- lifespan}{lifespan \times fraction \times\sqrt{2}} \right)]$ (Equation 11)

$erf(x) = \frac{2}{\sqrt{\pi}} \int_{0}^{x} e^{-t^{2}}dt$	(Equation 12)

where CDF is the cumulative distribution function and erf is the error function

As explained earlier, in *TimberTracer*, a portion of the HWPs decay undergoes recycling, another part is sent to landfills, and the remaining portion is combusted according to a predefined scheme. Given that each HWP has specific usage characteristics, the dynamic decay varies from one HWP to another, primarily influenced by lifespan and the recycling rate.

The following graphs showcase the decay of the initial carbon stock of HWPs among products with different lifespans (30, 15, and 5 years) without recycling (Left-side graph) and the decay of the initial stock considering various recycling rates (No recycling, 10%, and 20%) (Right-side graph).

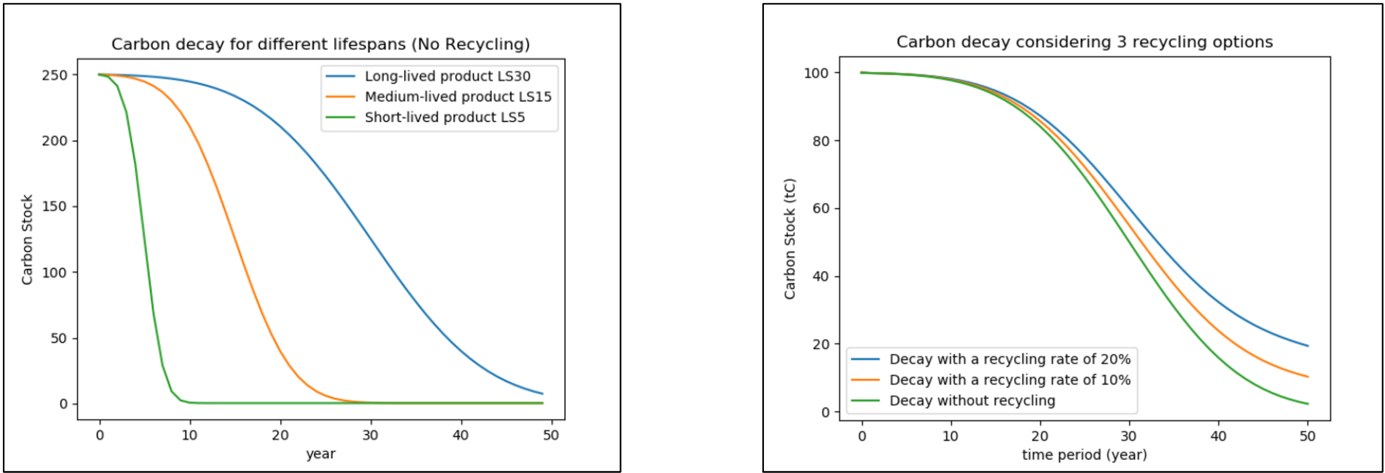

Figure 9 : Left side illustrates carbon stock decay among products with different lifespans (30, 15, and 5 years). The right side depicts carbon stock decay among stocks with different recycling rates (0%, 10%, and 20%).

These calculations are implemented by the model using the following functions:

**compute_R**

This function computes the annual recycling (for a period of n years) of a product, given its initial stock, lifespan, and recycling rate.

**compute_D**

This function computes the annual production of landfill or firewood proportionally to the annual decay of wood.

**compute_S**

This function computes the annual stock of a product given its initial stock, lifespan, and recycling rate.

**total_stock_calculator**

This function computes the total annual stock including the stock in HWPs and disposal sites.

**total_emission_calculator**

This function computes the total emissions from firewood, and disposal sites stock decomposition.

**total_recycling_calculator**

This function computes the total annual recycling of all the products given their use properties (i.e., lifespan and recycling rate).

The Table 6 below presents the structure of results rendered by the stock and flow module.

Table 6: An example of result structure from the stock and flow routine.

| **Year** | **Carbon stock** | **Carbon recycling** | **Carbon emissions** |
| --- | --- | --- | --- |
| 1 | - | - | - |
| . | - | - | - |
| . | - | - | - |
| N | - | - | - |

**5. The substitutor routine**

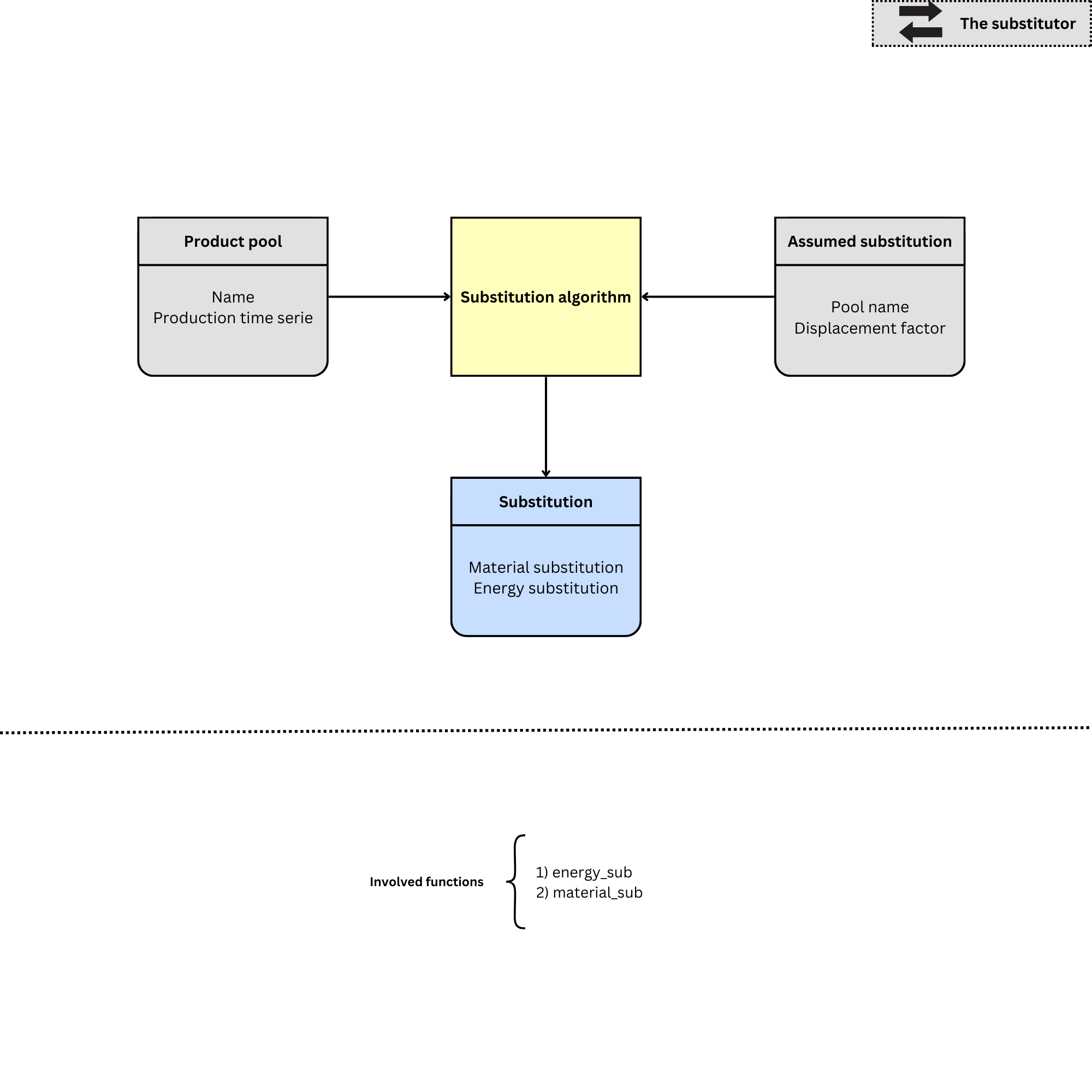

Figure 10: Graphical representation of the substitution routine

This module assesses the substitution effect resulting from the replacement of energy-intensive materials by wood products (material substitution) or the substitution of fossil fuel by bioenergy (energy substitution). This is practically achieved by multiplying the consumed HWP by a corresponding displacement factor defined as the avoided CO2 emissions corresponding to the use of a given dry volume of wood (m³). In the model, this computation is accomplished using two functions, namely **energy_sub** for energy substitution and **material_sub** for material substitution.

The Table 7 below presents the structure of results rendered by the substitution routine.

Table 7 : An example of result structure from the substitution routine.

| **Year** | **Energy substitution** | **Material substitution** |
| --- | --- | --- |
| 1 | - | - |
| . | - | - |
| . | - | - |
| N | - | - |
