## Supplementary material for "*TimberTracer*: A Comprehensive Framework for the Evaluation of Carbon Sequestration by Forest Management and Substitution of Harvested Wood Products": S4 (from another version of this preprint: version 3, 2025-01-15)

**S4: Data inputs of management and wood-use scenarios**

**Management scenarios inputs**

Table 1: Management inputs for the selective thinning scenario

| **N_rotation** | **age** | **total_pop** | **target_sum** | **DBH** | **bark** | **sap_area** | **heart_area** | **thinning_type** |
| --- | --- | --- | --- | --- | --- | --- | --- | --- |
| 1 | 39 | 1396 | 10.47 | 19.54 | 1.5 | 199.72 | 100.31 | "above" |
| 1 | 63 | 532 | 11.70 | 31.61 | 1.5 | 412.95 | 372.21 | "above" |
| 1 | 78 | 314 | 11.68 | 41.12 | 1.5 | 634 | 693 | "above" |
| 1 | 93 | 192 | 11.69 | 52.61 | 1.5 | 979 | 1195 | "above" |
| 1 | 108 | 123 | 11.53 | 65.3 | 1.5 | 1415 | 1933 | "above" |

Table 2: Management inputs for the clearcut scenario

| **N_rotation** | **age** | **total_pop** | **target_sum** | **DBH** | **bark** | **sap_area** | **heart_area** | **thinning_type** |
| --- | --- | --- | --- | --- | --- | --- | --- | --- |
| 1 | 39 | 1396 | 10.47 | 19.54 | 1.5 | 199.72 | 100.31 | "above" |
| 1 | 84 | 275 | 41.83 | 44.01 | 1.5 | 698.16 | 823.18 | "neutral" |

Table 3: Management inputs for the shelterwood scenario

| **N_rotation** | **age** | **total_pop** | **target_sum** | **DBH** | **bark** | **sap_area** | **heart_area** | **thinning_type** |
| --- | --- | --- | --- | --- | --- | --- | --- | --- |
| 1 | 39 | 1396 | 10.47 | 19.54 | 1.5 | 199.72 | 100.31 | "above" |
| 1 | 63 | 532 | 8.354 | 31.62 | 1.5 | 412.95 | 372.21 | "bottom" |
| 1 | 73 | 376 | 8.38 | 37.67 | 1.5 | 550.07 | 564.51 | "bottom" |
| 1 | 84 | 261 | 33.448 | 45.16 | 1.5 | 750.85 | 851.08 | "neutral" |
| 1 | 94 | 52 | 14.51 | 59.61 | 1.5 | 1552.41 | 1238.62 | "neutral" |

**Wood-use scenarios inputs**

Table 4: Wood-use inputs for the BAU scenario

| **product** | **r** | **ls** | **fire** | **landfill** |
| --- | --- | --- | --- | --- |
| furniture | 0.1 | 30 | 0.8 | 0.1 |
| lumber | 0.1 | 65 | 0.8 | 0.1 |
| paper | 0.1 | 1 | 0.8 | 0.1 |
| sawing | 0.1 | 50 | 0.8 | 0.1 |
| particle | 0.1 | 20 | 0.8 | 0.1 |

Table 5: Wood-use inputs for the reuse scenario

| **product** | **r** | **ls** | **fire** | **landfill** |
| --- | --- | --- | --- | --- |
| furniture | 0.11 | 30 | 0.79 | 0.1 |
| lumber | 0.11 | 65 | 0.79 | 0.1 |
| paper | 0.11 | 1 | 0.79 | 0.1 |
| sawing | 0.11 | 50 | 0.79 | 0.1 |
| particle | 0.11 | 20 | 0.79 | 0.1 |

Table 6: Wood-use inputs for the longevity scenario

| **product** | **r** | **Ls** | **fire** | **landfill** |
| --- | --- | --- | --- | --- |
| furniture | 0.1 | 33 | 0.8 | 0.1 |
| lumber | 0.1 | 71.5 | 0.8 | 0.1 |
| paper | 0.1 | 1.1 | 0.8 | 0.1 |
| sawing | 0.1 | 55 | 0.8 | 0.1 |
| particle | 0.1 | 22 | 0.8 | 0.1 |

Table 7: Wood-use inputs for the sustainability scenario

| **product** | **r** | **Ls** | **fire** | **landfill** |
| --- | --- | --- | --- | --- |
| furniture | 0,11 | 33 | 0,79 | 0,1 |
| lumber | 0,11 | 71.5 | 0,79 | 0,1 |
| paper | 0,11 | 1.1 | 0,79 | 0,1 |
| sawing | 0,11 | 55 | 0,79 | 0,1 |
| particle | 0,11 | 22 | 0,79 | 0,1 |

r, ls, fire, and landfill correspond to the recycling rate, lifespan, proportion of

decay allocated to fire, and proportion of decay that is landfilled, respectively.
