## Supplementary material for "*TimberTracer*: A Comprehensive Framework for the Evaluation of Carbon Sequestration by Forest Management and Substitution of Harvested Wood Products": S2 (from another version of this preprint: version 3, 2025-01-15)

**S2: Data on industrial processes, wood use, and substitution**

Table 1: Bucking allocation criteria according to different sectors (Cutting started from 30 cm above ground level)

| Product | Priority | Number | Length | Diameter | Quality criteria |
| --- | --- | --- | --- | --- | --- |
| Stump | 1 | 1 | 0.3 m |  | - |
| Interior carpentry | 2 | - | 3 – 5 m | Ø_SE_ ≥ 25 cm | (Ø_KW_ / Ø_HW_)^2^ ≤ 13% |
| Lumber | 3 | - | 3 – 6 m | Ø_SE_ ≥ 25 cm | Ø_KW_/ Ø_SE_ ≤ 30 % |
| Sawing | 4 | - | 3 – 5 m | Ø_SE_ ≥ 25 cm | - |
| Particle boards | 5 | 3 | 2 – 2.2 m | Ø_SE_ ≥ 7 cm | - |
| Paper pulp | 6 | - | 2.4 – 4.8 | Ø_SE_ ≥ 7 cm | - |
| Firewood | 7 | - | 0.5 – 1 m | Ø_SE_ ≥ 7 cm | - |
| Toplog | 8 | - | - |  | - |

Ø_SE_: Diameter small end, Ø_KW_: Diameter of knotted wood, Ø_HW_: Diameter of Heartwood, Ø_MED_: Median diameter

Table 2: Efficiency and loss of different wood products reported to the log volume including bark.

| **Product** | **Efficiency (%)** | **Loss (%)** |
| --- | --- | --- |
| Stump | 1 | 0 |
| Furniture | 0.45 | 0.55 |
| Lumber | 0.50 | 0.50 |
| Sawing | 0.40 | 0.60 |
| Particle boards | 0.76 | 0.24 |
| Paper pulp | 0.8 | 0.20 |
| Firewood | 1 | 0 |
| TopLog | 1 | 0 |

Table 3: Lifespan and recycling rate of different wood products and disposal.

| Wood product | Average lifespan (yrs.) | Recycling rate (%) |
| --- | --- | --- |
| Furniture | 30 | 0.1 |
| Lumber | 65 | 0.1 |
| Sawn wood | 50 | 0.1 |
| Particle boards | 20 | 0.1 |
| Paper pulp | 1 | 0.1 |
| Firewood including the BGB, stump, and the TopLog | 1 | - |
| Mill site dump | 5 | - |
| Landfill | 145 | - |

Table 4: Allocation of the harvested wood product losses.

|  | Fire (%) | Boards (%) | Paper (%) | Millsite (%) |
| --- | --- | --- | --- | --- |
| Furniture | 0.15 | 0.1 | 0.2 | 0.1 |
| Lumber | 0.15 | 0.1 | 0.15 | 0.1 |
| Sawing | 0.2 | 0.2 | 0 | 0.1 |
| Particle boards | 0.14 | 0 | 0 | 0.2 |
| Paper pulp | 0 | 0 | 0 | 0 |

Table 5: Assumed substitution between wood products and substitutes in the considered services. The fraction of product in the likely substitution scenario and the average reduction in CC impacts (aka. displacement factor) were documented from (Sathre & O’Connor, 2010; Suter et al., 2017).

| Wood product | Substitute | Service | Likely fraction of product in the substitution scenario (%) | The average reduction in CC impacts (tCO2-eq/m3) | Displacement factor (tCO2-eq/m3) |
| --- | --- | --- | --- | --- | --- |
| Furniture | Polypropylene | Furniture | 47 | 0.61 | 0.82 |
|  | Steel, chromium, secondary | Furniture | 47 | 0.99 |  |
|  | Steel, chromium, primary | Furniture | 6 | 1.23 |  |
| Lumber | Steel, secondary | Construction | 70 | -0.06 | 0.06 |
|  | Steel, primary | Construction | 10 | 0.58 |  |
|  | Concrete | Construction | 10 | 0.27 |  |
|  | Brick | Construction | 10 | 0.18 |  |
| Sawn wood | Concrete | Construction | 39 | 0.27 | 0.32 |
|  | Brick | Construction | 31 | 0.18 |  |
|  | Polyethylene | Packaging | 9 | 0.30 |  |
|  | Aluminum, secondary | Packaging | 0.9 | 0.09 |  |
|  | Aluminum, primary | Packaging | 0.1 | 3.67 |  |
|  | Polypropylene | Furniture | 10 | 0.30 |  |
|  | Steel, chromium, secondary | Furniture | 9 | 0.99 |  |
|  | Steel, chromium, primary | Furniture | 1 | 1.23 |  |
| Particle boards | Plaster | Construction | 60 | 0.06 | 0.2 |
|  | Glass | Furniture | 40 | 0.41 |  |
| Paper pulp | None | Print | 100 | - | - |
| Firewood including the TopLog | Natural gas | Energy | 30 | 0.32 | 0.48 |
|  | Fuel oil | Energy | 70 | 0.55 |  |

PS: The volume in the displacement factor refers to the dry volume.
