## Supplementary material for "*TimberTracer*: A Comprehensive Framework for the Evaluation of Carbon Sequestration by Forest Management and Substitution of Harvested Wood Products": S3 (from another version of this preprint: version 3, 2025-01-15)

**Table 1: Metadata of the TimberTracer’s data inputs and parameters**

| **Module** | **Generating function** | **Parameter name** | **Description** | **Values** | **Unit** | **Class** |
| --- | --- | --- | --- | --- | --- | --- |
| **Dendrometer** | dendrometer | scale | Weibull distribution scale parameter | - | - | numeric |
|  |  | shape | Weibull distribution shape parameter | - | - | numeric |
|  |  | total_population | Stand density | - | trees | integer |
|  |  | class_width | DBH class width | - | cm | integer |
|  |  | DBH | Stand diameter at breast height | - | cm | numeric |
|  | perform_thinning | tree_df | Stand dataframe | - | - | series |
|  |  | thinning_type | Type of the applied thinning | "above", "bottom", or "neutral" | - | character |
|  |  | target_sum | Basal area to be thinned | - | m2 | numeric |
|  |  | tolerance_g | Tolerance in reaching the recquired target_sum | - | m2 | numeric |
| **Bucking allocation** | taper_equation | a1, a2, b1, b2, b3, b4 | Taper equation parameters | - | - | numeric |
|  | knotted_wood_profile | a1, a2, b1, b2, b3, b4 | Same as described previously | - | - | - |
|  |  | a, c | Parameters of the height to the crown height model | - | - | numeric |
|  |  | alpha, beta | Parameters of the height to diameter model | - | - | numeric |
|  |  | ht | Total height of individual tree | - | m | numeric |
|  | sapwood_profile | a1, a2, b1, b2, b3, b4, ht, DBH | Same as described previously | - | - | - |
|  |  | sap_area | Sapwood area | - | cm2 | numeric |
|  |  | heart_area | Heartwood area | - | cm2 | numeric |
|  |  | dbh | DBH of individual tree | - | cm | numeric |
|  |  | bark | Bark thickness | - | cm | numeric |
|  | bucking_allocation | a1, a2, b1, b2, b3, b4, sap_area, heart_area, ht, DBH, a, c, alpha, beta, bark, dbh | Same parameters as described previously | - | - | - |
|  |  | tolerance_hfind | The tolerated deviation between the expected and the ratained ratio | - | - | numeric |
| **Processor** | processor | df | Dataframe of the thinned trees | - | - | series |
|  |  | efficiency | Effciency of the industrial processes of different logs | - | - | dictionary |
|  |  | loss_allocation | Reallocation of the industrials loss to different products | - | - | dictionary |
| **Stock and Flow** | compute_R | n | Number of years for the projection | - | years | integer |
|  |  | r | The eecycling rate of product | - | - | numeric |
|  |  | initial_stock | Initial carbon stock of product | - | tC | numeric |
|  |  | lifespan | Average lifetime of the product | - | years | numeric |
|  | compute_D | n, r, initial_stock, lifespan | Same parameters as described previously | - | - | - |
|  |  | k | The proportion of the decay to be allocated to a given pool | - | - | numeric |
|  | compute_S | n, r, initial_stock, lifespan | Same parameters as described previously | - | - | - |
|  | total_recycling_calculator | recycling_matrix | Recycling matrix holding recycing rates of all the products | - | - | dictionary |
|  |  | df, decay_matrix, n | Same parameters as described previously | - | - | - |
|  |  | K | Moisture content adjustment factor | - | - | numeric |
|  |  | D | Wood density | - | t/m3 | numeric |
|  |  | CC | Portion of carbon in dry biomass | - | tC/tDM | numeric |
|  | total_stock_calculator | df, recycling_matrix, decay_matrix, n, K, D, CC | Same parameters as described previously | - | - | - |
|  |  | t0.5 | Half-period of the degradation | - | years | numeric |
|  |  | L0 | Methane generation potential | - | tCH4/tWM | numeric |
|  |  | f | Fraction of methane in the released GHG | - | - | numeric |
|  |  | GWP_CH4 | Global warming potential of methane | - | CO2-eq | numeric |
|  | total_emission_calculator | df, recycling_matrix, decay_matrix, n, K, D, CC, t0.5, L0, f, GWP_CH4 | Same parameters as described previously | - | - | - |
| **Substitutor** | energy_sub | df, recycling_matrix, decay_matrix, n | Same parameters as described previously | - | - | - |
|  |  | substitution_matrix | Matrix of displacement factors | - | - | dictionary |
|  |  | C_CO2 | The conversion constant from C to CO2 | - | - | numeric |
|  | material_sub | df, recycling_matrix, decay_matrix, seubtitution_matrix, C_CO2, n | Same parameters as described previously | - | - | - |
| **Run** | run_model | scale, shape, class_width, a1, a2, b1, b2, b3, b4, a, c, alpha, beta, efficiency, loss_allocation, recycling_matrix, substitution_matrix, decay_matrix | Same parameters as described previously | - | - | - |
|  |  | tolerance_hfind | The tolerated deviation between the expected and the retained ratio corresponding to the search of a correspondant height | - | - | numeric |
|  |  | PH | Planning horizon | - | years | numeric |
|  |  | R | Management rotation | - | years | numeric |
|  |  | management_dic | Data on the management interventions | - | - | dictionary |
